# The Single-cell Pediatric Cancer Atlas: Data portal and open-source tools for single-cell transcriptomics of pediatric tumors

**DOI:** 10.1101/2024.04.19.590243

**Authors:** Allegra G. Hawkins, Joshua A. Shapiro, Stephanie J. Spielman, David S. Mejia, Deepashree Venkatesh Prasad, Nozomi Ichihara, Arkadii Yakovets, Avrohom M. Gottlieb, Kurt G. Wheeler, Chante J. Bethell, Steven M. Foltz, Jennifer O’Malley, Casey S. Greene, Jaclyn N. Taroni

## Abstract

The Single-cell Pediatric Cancer Atlas (ScPCA) Portal (https://scpca.alexslemonade.org/) is a data resource for uniformly processed single-cell and single-nuclei RNA sequencing (RNA-seq) data and de-identified metadata from pediatric tumor samples. Originally comprised of data from 10 projects funded by Alex’s Lemonade Stand Foundation (ALSF), the Portal currently contains summarized gene expression data for over 700 samples across 55 cancer types from ALSF-funded and community-contributed datasets. Downloads include gene expression data as `SinglecellExperiment` or `AnnData` objects containing raw and normalized counts, PCA and UMAP coordinates, and automated cell type annotations, along with summary reports. Some samples have additional data from bulk RNA-seq, spatial transcriptomics, and/or feature barcoding (e.g., CITE-seq and cell hashing) included in the download. All data on the Portal were uniformly processed using `scpca-nf`, an efficient and open-source Nextflow workflow that uses `alevin-fry` to quantify gene expression. Comprehensive documentation, including descriptions of file contents and a guide to getting started, is available at https://scpca.readthedocs.io.

## Introduction

The number of studies employing single-cell RNA-seq has grown rapidly since this technology was introduced [1]. Unlike its predecessor bulk RNA-seq, which averages the expression profiles of all cells within a sample, single-cell technology quantifies gene expression in individual cells. Tumors are known to be transcriptionally heterogeneous, which highlights the importance of using single-cell RNA-seq in studying tumor samples [2]. Researchers can use single-cell RNA-seq data of samples obtained from patient tumors to analyze and identify individual cell populations that may play important roles in tumor growth, resistance, and metastasis [3]. Additionally, single-cell RNA-seq data provides insight into how tumor cells interact with normal cells in the tumor microenvironment [4].

With the growing number of single-cell RNA-seq datasets, efforts have emerged to create centralized data resources. For example, resources like CELLxGENE [5,6] offer gene expression data from samples spanning hundreds of cell types in standardized analysis formats. Other resources offer harmonized data, which allows researchers to perform reliable cross-sample comparisons that leverage many biological contexts to complete their analysis and elucidate previously unknown similarities across samples and disease types. The Human Cell Atlas (HCA) and Human Tumor Atlas Network (HTAN) are two of many such resources. The HCA, which aims to use single-cell genomics to provide a comprehensive map of all cell types in the human body [7], contains uniformly processed single-cell RNA-seq data obtained from normal tissue with few samples derived from diseased tissue. The HTAN also hosts a collection of genomic data collected from tumors across multiple cancer types, including single-cell RNA-seq [8].

While existing resources have focused on making large quantities of harmonized data from normal tissue or adult tumor samples publicly available, there are considerably fewer efforts to harmonize and distribute data from pediatric tumors. Pediatric cancer is much less common than adult cancer, so the number of available samples from pediatric tumors is smaller compared to the number of adult tumors [9] and access to data from pediatric tumors is often limited. Thus, it is imperative to provide harmonized data from pediatric tumors to all pediatric cancer researchers [10]. To address this unmet need, Alex’s Lemonade Stand Foundation and the Childhood Cancer Data Lab developed and maintain the Single-cell Pediatric Cancer Atlas (ScPCA) Portal (https://scpca.alexslemonade.org/), an open-source data resource for single-cell and single-nuclei RNA-seq data of pediatric tumor samples.

The ScPCA Portal holds uniformly processed summarized gene expression from 10x Genomics droplet-based single-cell and single-nuclei RNA-seq for over 700 samples from a diverse set of 55 types of pediatric cancers. Originally comprised of data from 10 projects funded by Alex’s Lemonade Stand Foundation, the Portal has since expanded to include data contributed by pediatric cancer research community members. In addition to gene expression data from single-cell and single-nuclei RNA-seq, the Portal includes data obtained from bulk RNA-seq, spatial transcriptomics, and feature barcoding methods such as CITE-seq and cell hashing. All data on the Portal are available in formats ready for downstream analysis with common workflow ecosystems, such as `SinglecellExperiment` objects used by `R/Bioconductor` [11] or `AnnData` objects used by `Scanpy` and related Python modules [12]. Downloaded objects contain both raw and normalized gene expression counts, dimensionality reduction results, and cell type annotations.

To ensure that all current and future data on the Portal are uniformly processed, we created `scpca-nf`, an open-source Nextflow [13] pipeline (https://github.com/AlexsLemonade/scpca-nf). Using a consistent pipeline for all data increases transparency and allows users to perform analysis across multiple samples and projects without having to do any re-processing. The `scpca-nf` workflow uses `alevin-fry` [14] for fast and efficient quantification of single-cell gene expression for all samples on the Portal, including single-cell RNA-seq data and any associated CITE-seq or cell hash data. The `scpca-nf` pipeline also serves as a resource for the community, allowing others to process their own samples for comparison to samples available on the Portal and submit uniformly processed community contributions to the Portal.

Here, we present the Single-cell Pediatric Cancer Atlas as a freely available resource for all pediatric cancer researchers. The ScPCA Portal provides downloads ready for immediate use, allowing researchers to skip time-consuming data re-processing and wrangling steps. We provide comprehensive documentation about data processing and the contents of files on the Portal, including a guide to getting started working with an ScPCA dataset (https://scpca.readthedocs.io/). The ScPCA Portal advances pediatric cancer research by accelerating researchers’ ability to answer important biological questions.

## Results

### The Single-cell Pediatric Cancer Atlas Portal

In March of 2022, the Childhood Cancer Data Lab launched the Single-cell Pediatric Cancer Atlas (ScPCA) Portal to make uniformly processed, summarized single-cell and single-nuclei RNA-seq data and de-identified metadata from pediatric tumor samples openly available for download by the research community. Data available on the Portal was obtained using two different mechanisms: raw data was accepted from ALSF-funded investigators and processed using our open-source pipeline `scpca-nf`, or investigators processed their raw data using `scpca-nf` and submitted the output for inclusion on the Portal.

All samples on the Portal include a core set of metadata obtained from investigators, including age, sex, diagnosis, subdiagnosis (if applicable), tissue location, and disease stage. Some samples include additional metadata, such as treatment or tumor stage, if provided by the investigators. We standardized all provided metadata to maintain consistency across projects before adding it to the Portal. In addition to providing a human-readable value for the submitted metadata, we also provide ontology term identifiers, if applicable. Submitted metadata was mapped to associated ontology term identifiers obtained from HsapDv (age) [15], PATO (sex) [16,17], NCBI taxonomy (organism) [18,19], MONDO (disease) [20,21], UBERON (tissue) [22,23,24], and Hancestro (ethnicity, if applicable) [25,26]. These ontology term identifiers offer standardized metadata terms that facilitate comparisons among datasets within the Portal as well as to data from other research projects.

The Portal contains data from over 700 samples and 55 tumor types [27,28,29,30,31,32,33]. Figure 1A summarizes all samples from patient tumors and patient-derived xenografts currently available on the Portal. The total number of samples for each diagnosis is shown, along with the proportion of samples from each disease stage within a diagnosis group. The largest number of samples found on the Portal were obtained from patients with leukemia (n = 216). The Portal also includes samples from sarcoma and soft tissue tumors (n = 194), brain and central nervous system tumors (n = 167), and a variety of other solid tumors (n = 115). Most samples were collected at initial diagnosis (n = 520), with a smaller number of samples collected either at recurrence (n = 129), during progressive disease (n = 12), during or after treatment (n = 11), or post-mortem (n = 5). Along with the patient tumors, the Portal contains a small number of human tumor cell line samples (n = 6).

**Figure 1:**
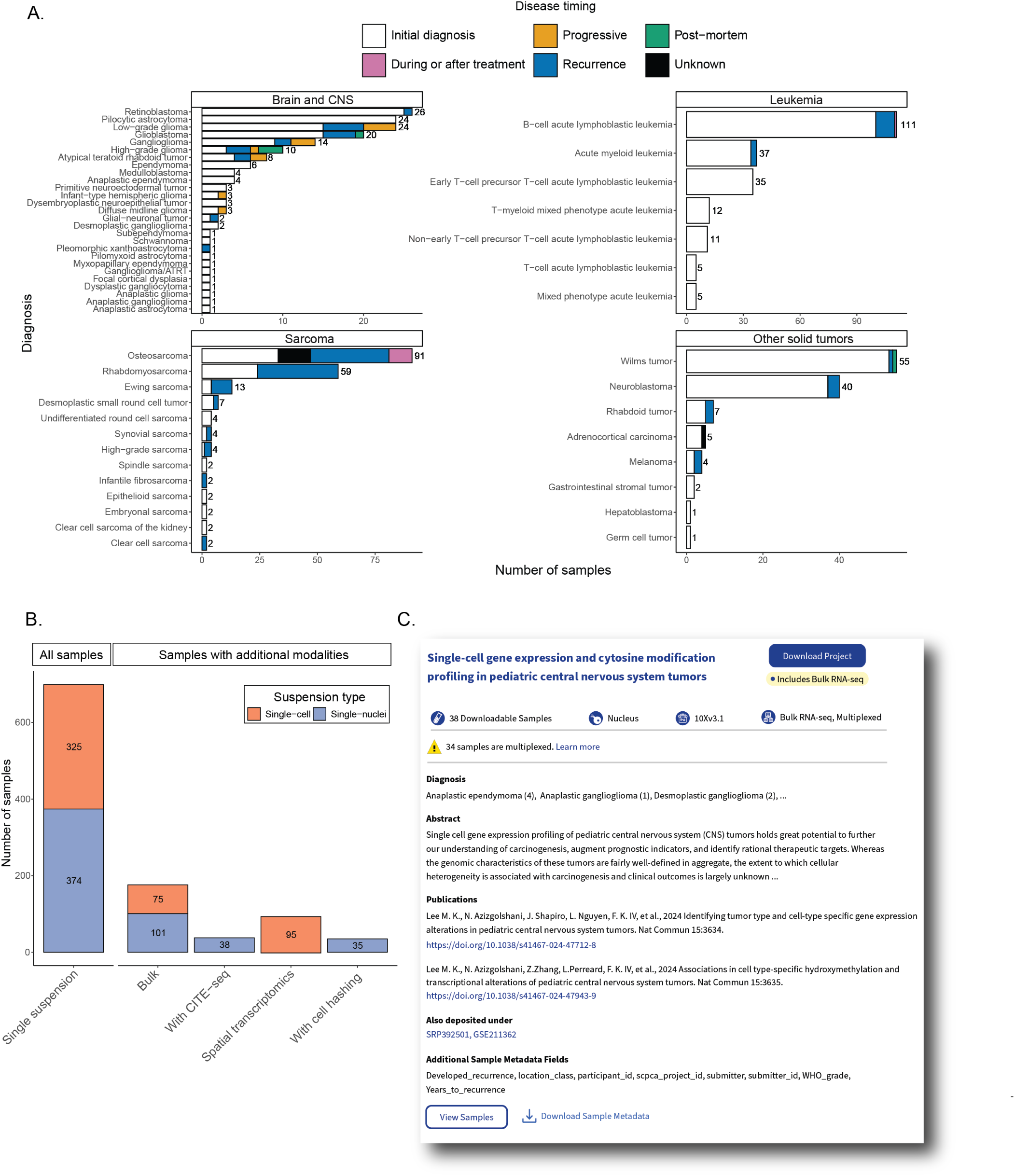
Overview of ScPCA Portal contents. A. Barplots showing sample counts across four main cancer groupings in the ScPCA Portal, with each bar displaying the number of samples for each cancer type. Each bar is colored based on the number of samples with the indicated disease timing, and total sample counts for each cancer type are shown to the right of each bar. B. Barplot showing sample counts across types of modalities present in the ScPCA Portal. All samples in the Portal are shown under the “All Samples” heading. Samples under the “Samples with additional modalities” heading represent a subset of the total samples with the given additional modality. Colors shown for each additional modality indicate the suspension type used, either single-cell or single-nuclei RNA-seq. For example, 75 single-cell samples and 101 single-nuclei samples have accompanying bulk RNA-seq data. Two samples were sequenced using both single-cell and single-nuclei suspensions so are included in the count for both “Single-cell” and “Single-nuclei” groups. Samples that were sequenced with either bulk RNA-seq or spatial transcriptomics and do not have accompanying single-cell or single-nuclei RNA-seq data are not represented in the total counts. C. Example of a project card as displayed on the “Browse” page of the ScPCA Portal. This project card is associated with project `ScPcP000006` [28,29]. Project cards include information about the number of samples, technologies and modalities, additional sample metadata information, submitter-provided diagnoses, and a submitter-provided abstract. Where available, submitter-provided citation information, as well as other databases where this data has been deposited, are also provided.

Each of the available samples contains summarized gene expression data from either single-cell or single-nuclei RNA sequencing. However, some samples also include additional data, such as CITE-seq quantification of cell-surface protein levels with antibody-derived tags (ADT) [34], or hashtag oligonucleotide (HTO) quantification for samples multiplexed prior to sequencing [35]. Out of the 704 samples, 95 have associated CITE-seq data, and 35 have associated multiplexing data. In some cases, multiple libraries from the same sample were collected for additional assays, either for bulk RNA-seq (n = 182) or spatial transcriptomics (n = 38). A summary of the number of samples with each additional modality is shown in Figure 1B, and a detailed summary of the total samples with each sequencing method broken down by project is available in Table S1.

Samples on the Portal are organized by project, where each project is a collection of similar samples from an individual lab. Users can filter projects based on diagnosis, included modalities (e.g., CITE-seq, bulk RNA-seq), 10x Genomics kit version (e.g., 10Xv2, 10Xv3), and whether or not a project includes samples derived from patient-derived xenografts or cell lines. The project card displays an abstract, the total number of samples included, a list of diagnoses for all samples included in the Project, and links to any external information associated with the project, such as publications and links to external data, such as SRA or GEO (Figure 1C). The project card also indicates the type(s) of sequencing performed, including the 10x Genomics kit version, the suspension type (cell or nucleus), if additional sequencing like bulk RNA-seq is present, or if the samples have been multiplexed using cell hashing.

### Uniform processing of data available on the ScPCA Portal

We developed `scpca-nf`, an open-source and efficient Nextflow [13] workflow for quantifying single-cell and single-nuclei RNA-seq data and processed all data available on the Portal with it. Using Nextflow as the backbone for the `scpca-nf` workflow ensures both reproducibility and portability.

All dependencies for the workflow are handled automatically, as each process in the workflow is run in a Docker container. Nextflow is compatible with various computing environments, including high-performance computing clusters and cloud-based computing, allowing users to run the workflow in their preferred environment. Setup requires organizing input files and updating a single configuration file for the computing environment after installing Nextflow and either Docker or Singularity. Nextflow will also handle parallelizing sample processing as allowed by the environment, minimizing run time. The combination of being able to execute a Nextflow workflow in any environment and run individual processes in Docker containers makes this workflow easily portable for external use.

When building `scpca-nf`, we sought a fast and memory-efficient tool for gene expression quantification to minimize processing costs. Due to its popularity, we expected many users of the Portal to process their own single-cell or single-nuclei data with Cell Ranger [36,37]. Thus, selecting a tool with comparable results to Cell Ranger was also desirable. In comparing `alevin-fry` [14] to Cell Ranger, we found `alevin-fry` had a lower run time and memory usage (Figure S1A), while retaining comparable mean gene expression for all genes (Figure S1B), total UMIs per cell (Figure S1C), and total genes detected per cell (Figure S1D). Based on these results, we used `salmon alevin` and `alevin-fry` [14] in `scpca-nf` to quantify gene expression data.

Taking FASTQ files as input, `scpca-nf` aligns reads using the selective alignment option in `salmon alevin` to an index with transcripts corresponding to spliced cDNA and intronic regions, denoted by `alevin-fry` as a `splici` index (Figure 2A). The output from `alevin-fry` includes a gene-by-cell count matrix for all barcodes identified, even those that may not contain true cells.

**Figure 2:**
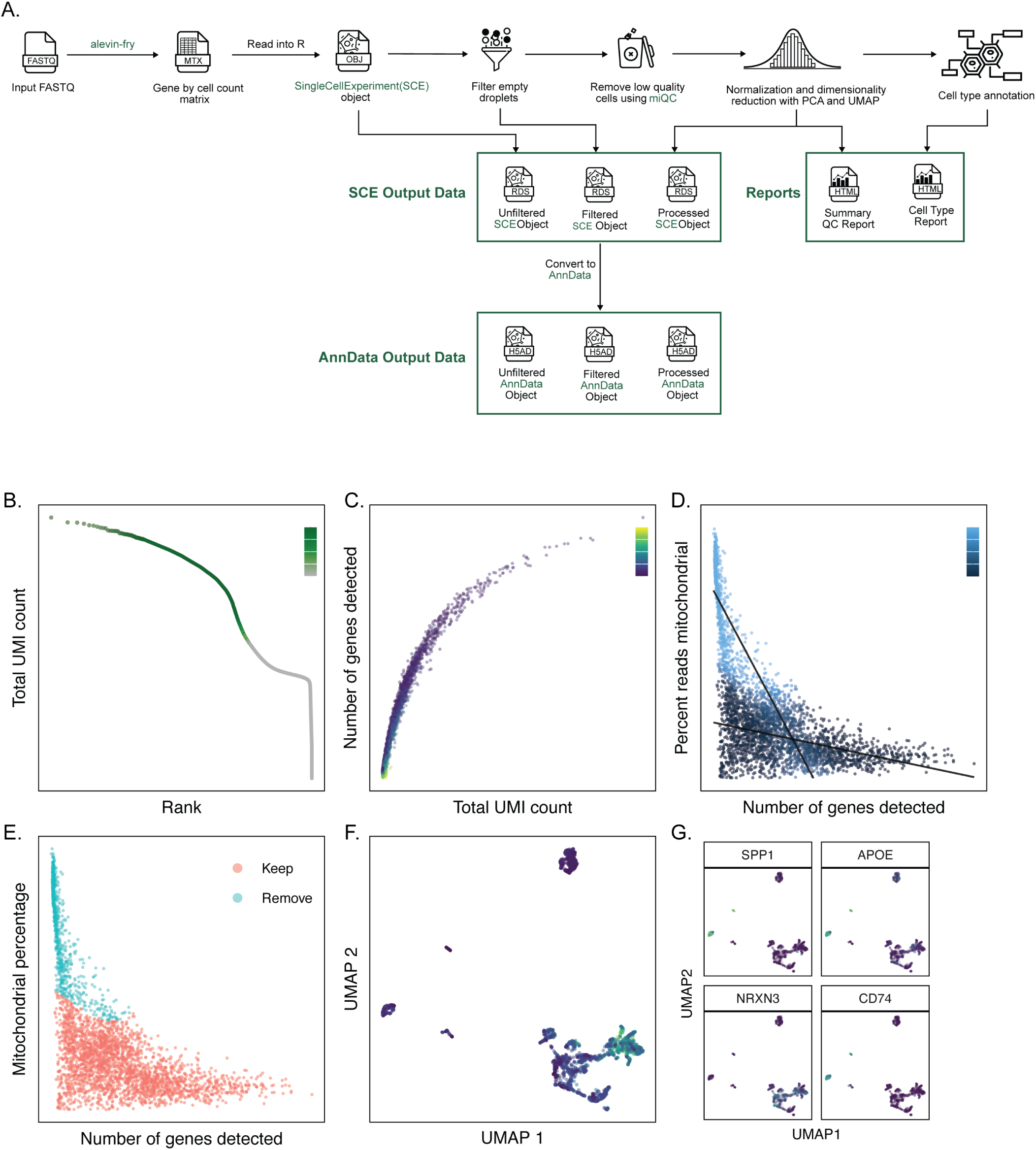
Overview of the scpca-nf workflow. A. Overview of `scpca-nf`, the primary workflow for processing single-cell and single-nuclei RNA-seq data for the ScPCA Portal. Mapping is first performed with `alevin-fry` to generate a gene-by-cell count matrix, which is read into `R` and converted into a `SinglecellExperiment` (`ScE`) object. This `ScE` object is exported as the `Unfiltered ScE Object` before further post-processing. Next, empty droplets are filtered out, and the resulting `ScE` is exported as the `Filtered ScE Object`. The filtered object undergoes additional post-processing, including removing low-quality cells, normalizing counts, and performing dimension reduction with principal components analysis and UMAP. The object undergoes cell type annotation and is exported as the `Processed ScE Object`. A summary QC report and a supplemental cell type report are prepared and exported. Finally, all `ScE` files are converted to `AnnData` format and exported. Panels B-G show example figures that appear in the summary QC report, shown here for `ScPcL000001` [32], as follows: The total UMI count for each cell in the `Unfiltered ScE Object`, ordered by rank. Points are colored by the percentage of cells that pass the empty droplets filter. B. The number of genes detected in each cell passing the empty droplets filter against the total UMI count. Points are colored by the percentage of mitochondrial reads in the cell. C. `miQc` model diagnostic plot showing the percent of mitochondrial reads in each cell against the number of genes detected in the `Filtered ScE Object`. Points are colored by the probability that the cell is compromised as determined by `miQc`. D. The percent of mitochondrial reads in each cell against the number of genes detected in each cell. Points are colored by whether the cell was kept or removed, as determined by both `miQc` and a minimum unique gene count cutoff, prior to normalization and dimensionality reduction. E. UMAP embeddings of log-normalized RNA expression values where each cell is colored by the number of genes detected. F. UMAP embeddings of log-normalized RNA expression values for the top four most variable genes, colored by the given gene’s expression. In the actual summary QC report, the top 12 most highly variable genes are shown.

`scpca-nf` performs filtering of empty droplets, removal of low-quality cells, normalization, dimensionality reduction, and cell type annotation (Figure 2A). The unfiltered gene-by-cell counts matrices are filtered to remove any barcodes that are not likely to contain cells using `DropletUtils::emptyDropscellRanger()` [38]. Low-quality cells are identified and removed with `miQc` [39], which jointly models the proportion of mitochondrial reads and detected genes per cell and calculates a probability that each cell is compromised. The remaining cells’ counts are normalized [40], and reduced-dimension representations are calculated using both principal component analysis (PCA) and uniform manifold approximation and projection (UMAP) [41]. Finally, cell types are classified using two automated methods, `SingleR` [42] and `cellAssign` [43].

To make downloading from the Portal convenient for R and Python users, downloads are available as either `SinglecellExperiment` or `AnnData` [44] objects. The workflow outputs a `SinglecellExperiment` object (saved as an `.rds` file) containing the fully processed results, including the dimension reduction results and cell type annotations, as well as objects containing the unfiltered and the empty droplet filtered gene-by-cell matrices. `scpca-nf` also converts all `SinglecellExperiment` objects to `AnnData` objects, which are saved as `.h5ad` files (Figure 2A). Downloads contain the unfiltered, filtered, and processed objects from `scpca-nf` to allow users to choose to perform their own filtering and normalization or to start their analysis from a processed object.

All downloads from the Portal include a quality control (QC) report with a summary of processing information (e.g., `alevin-fry` version), library statistics (e.g., the total number of cells), and a collection of diagnostic plots for each library (Figure 2B-G). A knee plot displaying total UMI counts for all droplets (i.e., including empty droplets) indicates the effects of the empty droplet filtering (Figure 2B). For each cell that remains after filtering empty droplets, the number of total UMIs, genes detected, and mitochondrial reads are calculated and summarized in a scatter plot (Figure 2C). We include plots showing the `miQc` model and which cells are kept and removed after filtering with `miQc` (Figure 2D-E). We also provide a UMAP plot with cells colored by the total number of genes detected and a faceted UMAP plot where cells are colored by the expression of a set of highly variable genes (Figure 2F-G).

### Processing samples with additional modalities

scpca-nf includes modules for processing samples with sequencing modalities beyond single-cell or single-nuclei RNA-seq data: corresponding ADT or CITE-seq data [34], multiplexed data via cell hashing [35], spatial transcriptomics, or bulk RNA-seq.

### Antibody-derived tags

To process ADT libraries, the ADT FASTQ files are quantified using `salmon alevin` and `alevin-fry` (Figure S2A). The output from `alevin-fry` is the unfiltered ADT-by-cell counts matrix. The ADT-by-cell counts matrix is read into R alongside the gene-by-cell counts matrix and saved as an alternative experiment (`altExp`) within the main `SinglecellExperiment` object containing the unfiltered RNA counts. The workflow performs ADT-by-cell counts matrix normalization (see Methods for details), but no filtering based on ADT expression or quality is performed. Instead, we report QC statistics that users can employ for additional filtering before performing downstream analyses.

If a library contains ADT data, the QC report will include an additional section with a summary of ADT-related statistics, such as how many cells express each ADT, and ADT-specific diagnostic plots (Figure S2B-D). We include plots summarizing the potential effects of removing low-quality cells based on RNA and ADT counts in the QC report (Figure S2B). The first quadrant indicates which cells would be kept if the object were filtered using both RNA and ADT quality measures. The other facets highlight which cells would be removed if filtering were done using only RNA counts, only ADT counts, or both. The top four ADTs with the most variable expression are also identified and visualized using density plots to show the normalized ADT expression across all cells (Figure S2C) and UMAPs – calculated from RNA expression data – with cells colored by ADT expression (Figure S2D).

### Multiplexed libraries

To process multiplexed libraries, the HTO FASTQ files are quantified using `salmon alevin` and `alevin-fry` (Figure S2C). As with ADT data, the HTO-by-cell counts matrix produced by `alevin-fry` is saved as an `altExp` within the main `SinglecellExperiment` object.

Although `scpca-nf` quantifies the HTO data and includes an HTO-by-cell counts matrix in all objects, `scpca-nf` does not demultiplex the samples into one sample per library. Instead, `scpca-nf` applies multiple demultiplexing methods, including demultiplexing with `DropletUtils::hashedDrops()` [45], demultiplexing with `Seurat::HTODemux()` [35], and genetic demultiplexing when bulk RNA-seq data are available. `scpca-nf` uses the genetic demultiplexing method described in Weber et al. [46], which uses bulk RNA-seq as a reference for the expected genotypes found in each single-cell RNA-seq sample. The results from all available demultiplexing methods are saved in the filtered and processed `SinglecellExperiment` objects.

If a library has associated HTO data, an additional section is included in the `scpca-nf` QC report. This section summarizes HTO-specific library statistics, such as how many cells express each HTO. No additional plots are produced, but a table summarizing the results from all three demultiplexing methods is included.

### Bulk and spatial transcriptomics

Some samples also include data from bulk RNA-seq and/or spatial transcriptomics libraries. Both of these additional sequencing methods are supported by `scpca-nf`. To quantify bulk RNA-seq data, `scpca-nf` takes bulk FASTQ files as input, trims reads using `fastp` [47], and then aligns and quantifies reads with `salmon` (Figure S3A) [48]. The output is a single TSV file with the gene-by-sample counts matrix for all samples in a given ScPCA project.

To quantify spatial transcriptomics data, `scpca-nf` takes the RNA FASTQ and slide image as input (Figure S3B). As `alevin-fry` does not yet fully support spatial transcriptomics data, `scpca-nf` uses Space Ranger to quantify all spatial transcriptomics data [49]. The output includes the spot-by-gene matrix along with a summary report produced by Space Ranger.

### Downloading projects from the ScPCA Portal

On the Portal, users can either download data from individual samples or all data from an entire ScPCA project. When downloading data for an entire project, users can choose between receiving the individual files for each sample (default) or one file containing the gene expression data and metadata for all samples in the project as a merged object. Users also have the option to choose their desired format and receive the data as `SinglecellExperiment` (`.rds`) or `AnnData` (`.h5ad`) objects.

For downloads with samples as individual files, the download folder will include a sub-folder for each sample in the project (Figure 3A). Each sample folder contains all three object types (unfiltered, filtered, and processed) in the requested file format and the QC and cell type summary report for all libraries from the given sample. The objects house the summarized gene expression data and associated metadata for the library indicated in the filename.

**Figure 3:**
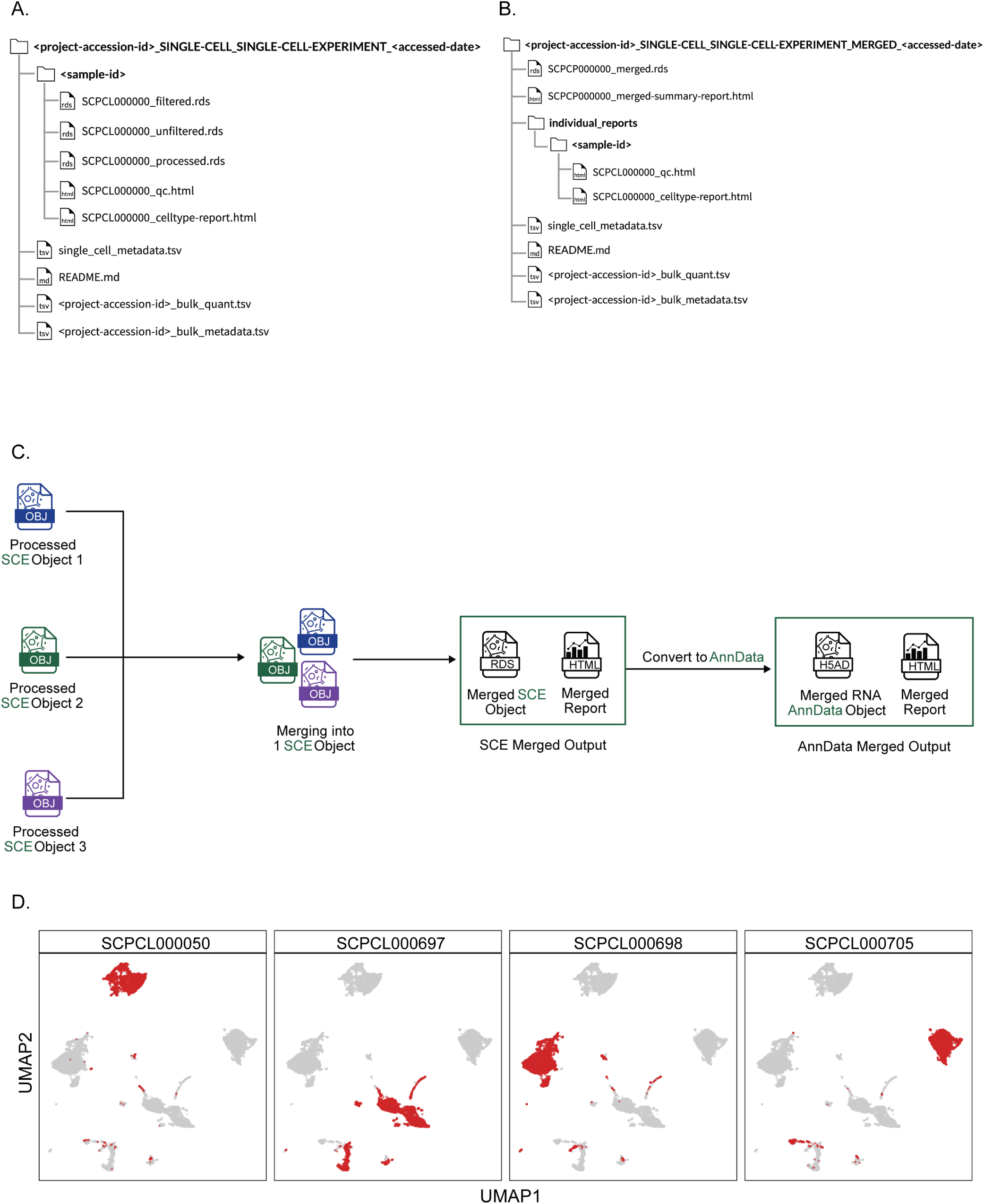
ScPCA Portal project download file structure and merged object workflow. A. File download structure for an ScPCA Portal project download in `SinglecellExperiment` (`ScE`) format. The download folder is named according to the project ID, data format, and the date it was downloaded. Download folders contain one folder for each sample ID, each containing the three versions (unfiltered, filtered, and processed) of the expression data as well as the summary QC report and cell type report all named according to the ScPCA library ID. The `single_cell_metadata.tsv` file contains sample metadata for all samples included in the download. The `README.md` file provides information about the contents of each download file, additional contact and citation information, and terms of use for data downloaded from the ScPCA Portal. The files `_bulk_quant.tsv` and `_bulk_metadata.tsv` are only present for projects that also have bulk RNA-Seq data and contain, respectively, a gene-by-sample matrix of raw gene expression as quantified by `salmon`, and associated metadata for all samples with bulk RNA-Seq data. B. File download structure for an ScPCA Portal merged project download in `ScE` format. The download folder is named according to the project ID, data format, and the date it was downloaded. Download folders contain a single merged object containing all samples in the given project as well as a summary report briefly detailing the contents of the merged object. All summary QC and cell type reports for each individual library are also provided in the `individual_reports` folder arranged by their sample ID. As in panel (A), additional files `single_cell_metadata.tsv`, `_bulk_quant.tsv`, `_bulk_metadata.tsv`, and `README.md` are also included. C. Overview of the merged workflow. Processed `ScE` objects associated with a given project are merged into a single object, including ADT counts from CITE-seq data if present, and a merged summary report is generated. Merged objects are available for download either in `ScE` or `AnnData` format. D. Example of UMAPs as shown in the merged summary report. A grid of UMAPs is shown for each library in the merged object, with cells in the library of interest shown in red and cells belonging to other libraries shown in gray. The UMAP is constructed from the merged object such that all libraries contribute an equal weight, but no batch correction was performed. The libraries pictured are a subset of libraries in the ScPCA project `ScPcP000003`. For this figure specifically, the merged UMAP was constructed from a merged object containing only these four libraries, but the merged object and summary report on the ScPCA Portal for `ScPcP000003` contain all of this project’s libraries.

All project downloads include a metadata file, `single_cell_metadata.tsv`, containing relevant metadata for all samples, and a `README.md` with information about the contents of each download, contact and citation information, and terms of use for data downloaded from the Portal (Figure 3A-B). If the ScPCA project includes samples with bulk RNA-seq, two additional files are included: a gene-by-sample counts matrix (`_bulk_quant.tsv`) with the quantified gene expression data for all samples in the project, and a metadata file (`_bulk_metadata.tsv`).

### Merged objects

Providing data for all samples within a single file facilitates performing joint gene-level analyses, such as differential expression or gene set enrichment analyses, on multiple samples simultaneously.

Therefore, we provide a single, merged object for each project containing all raw and normalized gene expression data and metadata for all single-cell and single-nuclei RNA-seq libraries within a given ScPCA project (with some exceptions as described in the Methods). If downloading data from an ScPCA project as a single, merged file, the download will include a single `.rds` or `.h5ad` file, a summary report for the merged object, and a folder with all individual QC and cell type reports for each library found in the merged object (Figure 3B).

To build the merged objects, we created an additional stand-alone workflow for merging the output from `scpca-nf`, `merge.nf` (Figure 3C). `merge.nf` takes the processed `SinglecellExperiment` objects for all single-cell and single-nuclei libraries in a given ScPCA project as input and produces a single merged gene-by-cell counts matrix containing all cells from all libraries. No batch correction or integration is performed when creating the merged object. Where possible, library-, cell- and gene-specific metadata found in the individual processed `SinglecellExperiment` objects are also merged. The merged normalized counts matrix is then used to select high-variance genes in a library-aware manner before performing dimensionality reduction with both PCA and UMAP. If additional modalities are present, these are similarly merged and included in the output object (see Methods). `merge.nf` outputs the merged and processed object as a `SinglecellExperiment` object. All merged `SinglecellExperiment` objects are converted to `AnnData` objects and exported as `.h5ad` files.

`merge.nf` outputs a summary report for each merged object, which includes a set of tables summarizing the types of samples and libraries included in the project, such as types of diagnosis, and a faceted UMAP showing all cells from all libraries. Figure 3D shows an example of this plot with a subset of libraries from an ScPCA project.

### Annotating cell types

Assigning cell type labels to single-cell and single-nuclei RNA-seq data is often an essential step in analysis. Cell type annotation requires knowledge of the expected cell types in a dataset and associated gene expression patterns for each cell type, which may be available in other public databases or individual publications. Automated cell type annotation methods leveraging public databases are an excellent initial step in the labeling process, as they can be applied consistently and transparently across all samples in a dataset. As such, we include cell type annotations determined using two different automated methods, `SingleR` [42] and `cellAssign` [43], in all processed `SinglecellExperiment` and `AnnData` objects.

Most public annotated reference datasets that can be used with `SingleR` and `cellAssign` – including those we use for the Portal – are derived from normal tissue, making accurately annotating tumor datasets particularly difficult. Observing consistent cell type annotations across methods can indicate higher confidence in the provided labels, so we created a set of ontology-aware rules to assign consensus cell type labels based on the methods’ agreement. These consensus cell type assignments can be found in all processed `SinglecellExperiment` and `AnnData` objects on the Portal.

For some ScPCA projects, submitters provided their own curated cell type annotations, including annotation of tumor cells and disease-specific cell states. These submitter-provided annotations can be found in all `SinglecellExperiment` and `AnnData` objects (unfiltered, filtered, and processed).

### Choosing cell typing references

`SingleR` is a reference-based annotation method that requires an existing bulk or single-cell RNA-seq dataset with annotations. To identify an appropriate reference to use with `SingleR`, we annotated a small number of samples across multiple disease types with all human-specific references available in the `celldex` package [42]. The output from `SingleR` includes a score matrix containing a score for each cell and all possible cell types found in the reference, where higher scores are associated with assigned cell types. We calculated the delta median statistic for each cell in the dataset by subtracting the median score from the score associated with the assigned cell type label. The delta median statistic helps evaluate how confident `SingleR` is in assigning each cell to a specific cell type, where low delta median values indicate ambiguous assignments and high delta median values indicate confident assignments [50]. This measure showed that the `BlueprintEncodeData` reference [51,52], which includes a variety of normal cell types, performed similarly to or better than other references when applied to samples from a variety of diagnoses (Figure S4). Based on these findings, we used the `BlueprintEncodeData` reference to annotate cells from all libraries on the Portal. Use of a consistent reference also supports cross-project analyses.

In contrast, `cellAssign` is a marker-gene-based annotation method that requires a binary matrix with all cell types and all associated marker genes as the reference. We used the list of marker genes available as part of `PanglaoDB` [53] to construct organ-specific marker gene matrices with marker genes from all cell types listed for the specified organ. Since many cancers may have infiltrating immune cells, all immune cells were also included in each organ-specific reference. For each ScPCA project, we used the organ-specific marker gene matrix that most closely matched the tissue type from which the sample was obtained (e.g., for brain tumors, we used a brain-specific marker gene matrix with all brain and immune cell types). If `cellAssign` cannot find a likely cell type from the marker gene matrix, it does not assign a cell type. Because we annotate cells from tumor samples using references containing only normal cells, we anticipate that many cells, particularly the tumor cells, will not have a suitable cell type match in the reference. Indeed, when applying `cellAssign` to tumor samples with our chosen reference, we observed that many cells were labeled as `Unknown` (Figure S5A). When comparing annotations obtained from `cellAssign` and `SingleR` annotations to submitter-provided annotations, we noticed the labels for non-tumor cells were similar between `cellAssign`, `SingleR`, and submitter annotations, while the tumor cells were not assigned using `cellAssign` (Figure S5B).

### Adding cell type annotations to the ScPCA Portal

`scpca-nf` adds cell type annotations from `SingleR` and `cellAssign` to all processed `SinglecellExperiment` objects (Figure 4A). This requires two additional reference files as input to the workflow: a classification model built from a reference dataset for `SingleR` and a marker gene matrix for `cellAssign`. We include assigned cell type labels and additional output from each method – the `SingleR` score matrix and the `cellAssign` prediction matrix, which contains a probability that a cell is of a cell type – to each processed object available from the Portal.

**Figure 4:**
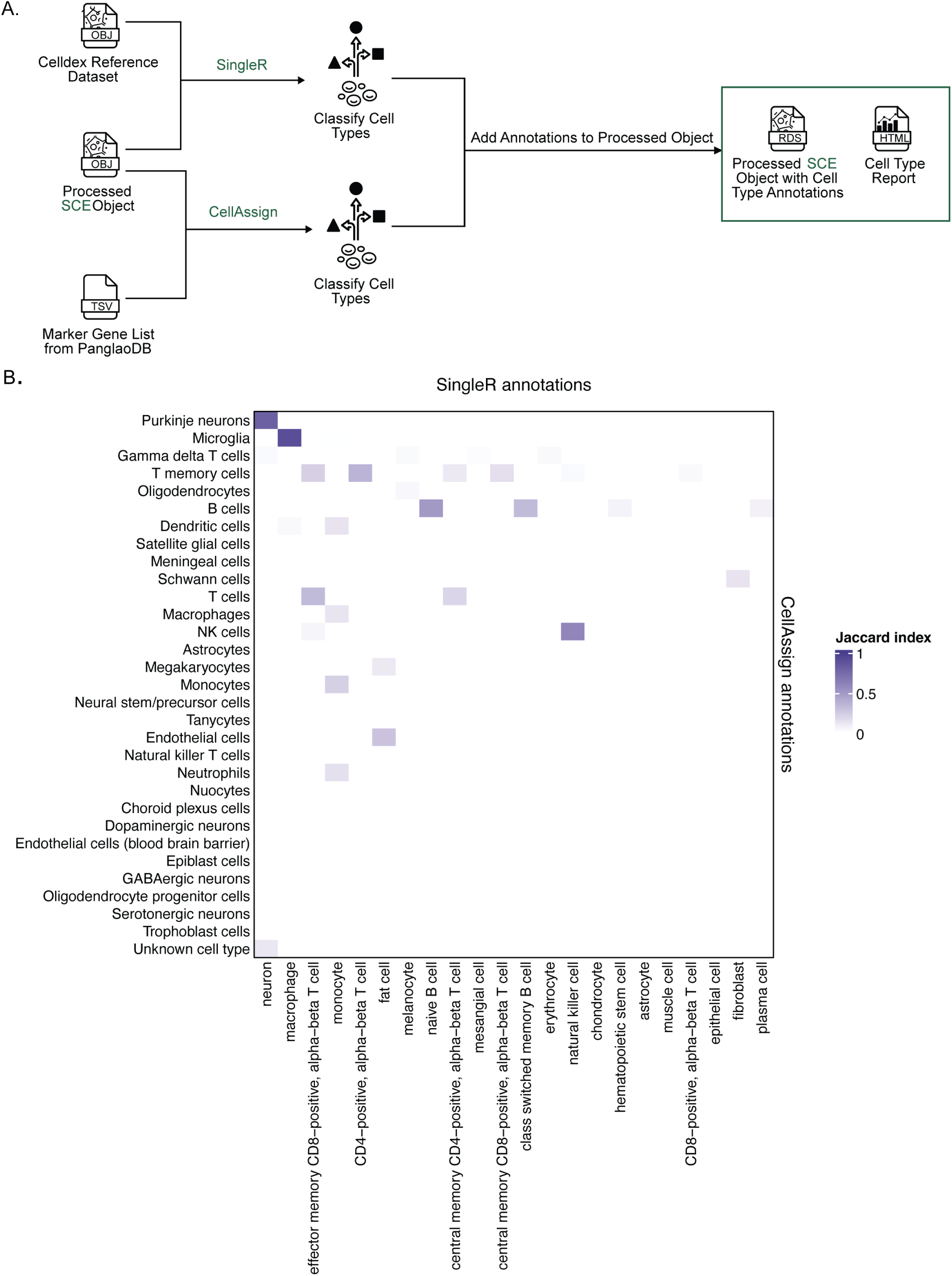
Cell type annotation in scpca-nf. A. Expanded view of the process for adding cell type annotations within `scpca-nf`, as introduced in Figure 2A. Cell type annotation is performed on the `Processed ScE Object`. A `celldex` [42] reference dataset with ontology labels is used as input for annotation with `SingleR` [42], and a list of marker genes compiled from `PanglaoDB` [53] is used as input for annotation with `cellAssign` [43]. Results from these automated cell type annotation methods and a consensus cell type annotation are then added to the `Processed ScE Object`. A cell type summary report with information about reference sources, comparisons among cell type annotation methods, and diagnostic plots is created. Although not shown in this panel, cell type annotations are also included in the `Processed AnnData Object` created from the `Processed ScE Object` (Figure 2A). B. Example heatmap as shown in the cell type summary report comparing annotations with `SingleR` and `cellAssign`. Heatmap cells are colored by the Jaccard similarity index. A value of 1 means that there is complete overlap between which cells are annotated with the two labels being compared, and a value of 0 means that there is no overlap between which cells are annotated with the two labels being compared. The heatmap shown is from library `ScPcL000468` [27].

We also produce an additional cell type report with information about reference sources, comparisons among cell type annotation methods, and diagnostic plots. Tables summarizing the number of cells assigned to each cell type for each method are shown alongside UMAPs coloring cells by the assigned cell type. We calculate the Jaccard index between pairs of cell type labels to compare annotations between the two methods and display it in a heatmap (an example is shown in Figure 4B). Jaccard index values close to 1 indicate high agreement and a high proportion of overlapping cells, which may indicate higher confidence predictions.

The report also includes diagnostic plots for each method. To evaluate confidence in `SingleR` cell type annotations, the delta median statistic is calculated by subtracting the median score from the score associated with the assigned cell type label [50]. The cell type report shows the distribution of delta median values for each cell type. A higher delta median statistic for a cell generally indicates higher confidence in the final cell type annotation. We also display the distribution of all probabilities calculated by `cellAssign`; more confident labels are expected to have many values close to 1.

If the submitter provided cell type annotations, the cell type report also includes a table summarizing the submitter cell type annotations, a UMAP plot in which each cell is colored by the submitter annotation, and a comparison of the submitter annotations to the automated cell typing results from `SingleR` and `cellAssign`. The Jaccard index is calculated for all pairs of cell type labels in submitter annotations and `SingleR` annotations, and in submitter annotations and `cellAssign` annotations. The results from both comparisons are displayed in a stacked heatmap available in the report, an example of which is shown in Figure S5B.

### Assigning consensus cell types

`SingleR` and `cellAssign` use different references and distinct computational approaches to label cells. We expect cells with the same or similar cell type labels from both methods to be more accurately annotated. `scpca-nf` therefore assigns consensus cell type labels when the two automated methods agree. To account for different levels of granularity in reference datasets, we employed an ontology-based approach to assign a consensus cell type label. Specifically, the consensus cell type annotation is equivalent to the latest common ancestor (LCA) in Cell Ontology [54,55,doi? 10.1186/gb-2005-6-2-r21] shared between the two predicted cell types. To ensure specificity in the consensus labels, cells were only assigned a consensus cell type if the identified LCA had no more than 170 descendant terms, with a few exceptions (see Methods for more details). We chose this threshold to exclude overly general cell ontology terms, such as lymphocyte, while retaining meaningful classifications like T cell and B cell. After assigning all consensus cell types, we looked at the expression of cell-type-specific marker genes across all cells to validate the assignments (Figure 5A, Figure S6).

**Figure 5:**
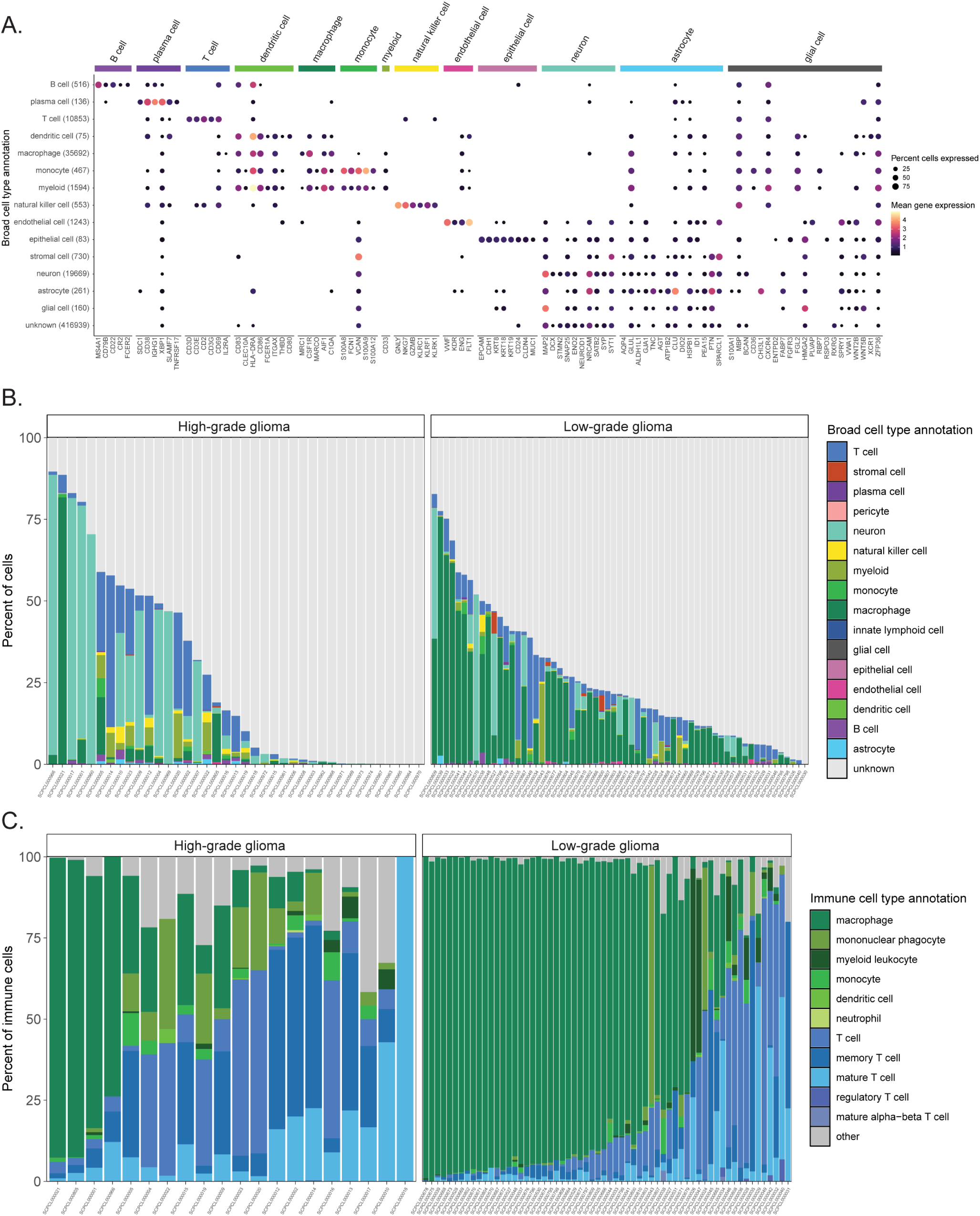
Consensus cell type annotations in Brain and CNS tumors. A. Dot plot showing expression of cell-type-specific marker genes across all libraries from brain and central nervous system (CNS) tumors, excluding multiplexed libraries. Expression is shown for each broad cell type annotation, where each broad cell type annotation is a collection of similar consensus cell type annotations. The y-axis displays the broad consensus cell type observed across libraries, with the total number of cells indicated in parentheses. The x-axis displays marker genes, determined by `cellMarker2.0` [72], used for consensus cell type validation for each cell type shown along the top annotation bar. Dots are colored by mean gene expression across libraries and sized proportionally to the percent of libraries they are observed in, out of all cells with the same broad cell type annotation in brain and CNS tumor libraries. B. Barplot showing the percentage of each broad consensus cell type annotation across libraries of brain and CNS tumors, separated into high-grade (left panel) and low-grade (right panel) glioma diagnoses for non-multiplexed libraries. C. Barplot showing all consensus cell types classified as immune cells across libraries of brain and CNS tumors, separated into high-grade (left panel) and low-grade (right panel) glioma diagnoses for non-multiplexed libraries. The percentage shown corresponds to the percentage of immune cells classified as the indicated consensus cell type. Only libraries comprised of at least 1% immune cells, based on consensus cell type annotations, are shown. Specific consensus cell types for myeloid and lymphocyte immune cells are shown, with all other consensus immune cell types included in “other.” Notably, granulocytes are also included in “other” because only 1 granulocyte was present in all libraries shown (specifically, `ScPcL000763`).

The consensus cell type labels provide harmonized cell type annotations for all samples in the ScPCA Portal, facilitating downstream analyses across multiple samples. Consensus annotations can be particularly useful when examining samples from multiple projects submitted by different investigators. For example, we show the distribution of cell types observed in all high-grade and low-grade glioma samples in Figure 5B, which originate from six different projects and four different investigators. Here, we can identify similar cell types across all glioma samples, but the composition of cell types present in each sample is heterogeneous.

Previous studies have characterized the glioma immune microenvironment as being predominantly composed of myeloid cells, including microglia and glioma-associated macrophages, with smaller proportions of lymphocytes such as T cells [56,57]. Focusing on the immune infiltrate in glioma samples reveals that most immune cells in ScPCA samples are classified as either myeloid or T cell types. However, there is notable heterogeneity even within HGG and LGG subtypes (Figure 5C). A summary of all the consensus cell types observed in all other ScPCA samples can be found in Figure S7.

### Analysis of bulk RNA-seq

Several projects in the ScPCA Portal contain bulk RNA-seq data in addition to single-cell/nuclei RNA-seq data. Previous research has suggested that, compared to bulk RNA-seq, single-cell/nuclei RNA-seq technologies may fail to capture certain cell types [58], for example, due to technical aspects of library preparation. We therefore asked whether we could identify differences in biological signal between these two modalities that may suggest distinct cell type distributions. We specifically focused on ScPCA projects with solid tumors, considering only samples with both sequencing modalities, and excluded low-quality single-cell/nuclei libraries and multiplexed samples. We analyzed 97 samples across five projects: `ScPcP00001`, `ScPcP000002`, `ScPcP000006`, `ScPcP000006`, and `ScPcP000017`. Projects `ScPcP000001` and `ScPcP000002` comprise high- and low-grade gliomas, respectively, and were sequenced at the bulk and single-cell levels. `ScPcP000006`, `ScPcP000006`, and `ScPcP000017` comprise Wilms tumors, CNS tumors, and osteosarcomas, respectively, and were sequenced at the bulk and single-nuclei levels. As described in the Methods, we derived pseudobulk expression matrices for each single-cell/nuclei library, and we compared their expression to bulk using a series of linear models (one per ScPCA project) predicting bulk from pseudobulk expression with a random effect controlling for sample (Figure 6A, Figure S8A). Across all projects, we observed a positive relationship between bulk and pseudobulk expression, consistent with our expectations.

**Figure 6:**
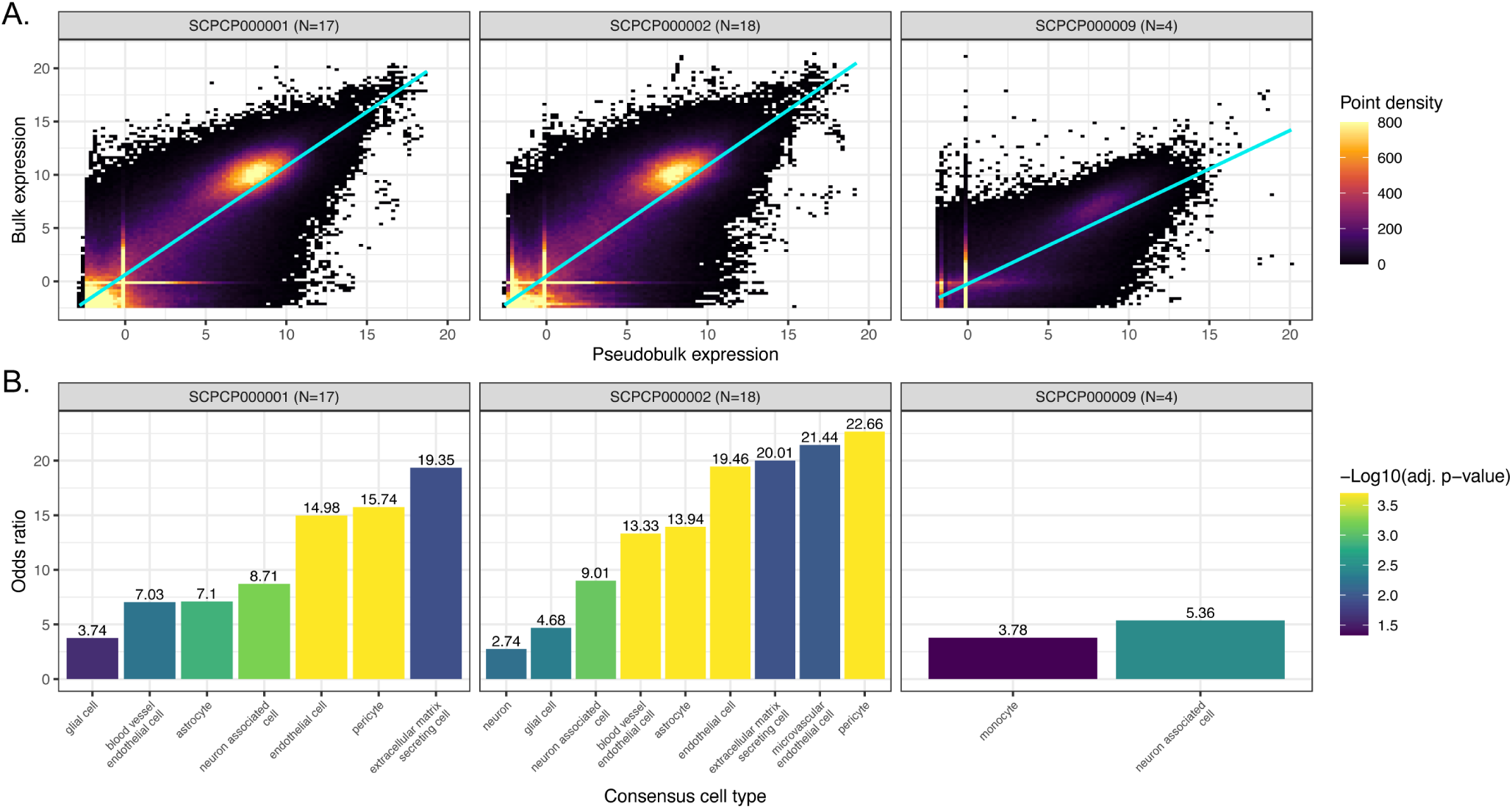
Comparison of bulk and pseudobulk modalities. A. Scatter plots colored by point density of `DESeq2` -transformed and normalized bulk RNA-seq expression compared to pseudobulk expression from single-cell/nuclei RNA-seq. Samples with RNA-seq for both bulk and single-cell/nuclei modalities, excluding multiplexed samples, from ScPCA projects comprising brain and central nervous system tumors are shown, with the number of samples considered per project shown in parentheses. The regression line is also shown for each project. Results from additional projects are shown in Figure S8A. B. Odds ratios from overrepresentation analysis for the same samples shown in panel A, colored by FDR-corrected significance. Each odds ratio represents the odds that marker genes for the given cell type were overrepresented in bulk RNA-seq when compared to single-cell/nuclei RNA-seq, relative to other genes. A total of 36 consensus cell types were evaluated for each project shown here. Results from additional projects are shown in Figure S8B.

We next performed an overrepresentation analysis to probe for differences in gene expression that might suggest differences in cell type composition and/or abundance between modalities. To this end, we calculated the per-gene median of each project’s model residuals and identified outliers, where “positive outliers” are genes with higher bulk RNA-seq expression than expected from pseudobulk expression, and conversely “negative outliers” are genes with lower bulk RNA-seq expression than expected from pseudobulk expression. Using marker gene sets associated with consensus cell types, we calculated the odds ratio in each direction as the odds a cell type marker gene is present in the given outlier direction compared to other genes. Following permutation testing and P-value correction to control the FDR at 5%, we indeed found that several cell type marker gene sets had higher, but never lower, bulk RNA-seq expression than expected (Figure 6B, Figure S8B).

In brain and CNS tumors, the marker genes overrepresented in bulk RNA-seq expression corresponded nearly exclusively to stromal (e.g., endothelial and extracellular matrix secreting cells) and/or neuronal cell types (e.g., glial cells and astrocytes), all of which are known to be prevalent non-immune cells in glioma tumor microenvironments [59,60] (Figure 6B). The only exceptions were monocyte marker genes being overrepresented in bulk RNA-seq expression for `ScPcP000006` (brain and CNS tumors), which was sequenced at the single-nuclei level, while projects `ScPcP000001` (high-grade gliomas) and `ScPcP000002` (low-grade gliomas) were sequenced at the single-cell level. This difference may reflect the increased sensitivity of single-cell approaches to detecting immune cells relative to single-nuclei approaches [61].

Given that our consensus cell type analysis identified various immune cells from high- and low-grade gliomas (Figure 5), these results suggest that non-immune cells may have been lost during single-cell library preparation. Indeed, several of these overrepresented bulk cell types for `ScPcP000001` and `ScPcP000002` were not among the single-cell consensus cell types themselves (`ScPcP000001` : “blood vessel endothelial cell”, “extracellular matrix secreting cell”, “glial cell”, “pericyte”; `ScPcP000002` : “blood vessel endothelial cell”, “extracellular matrix secreting cell”, “microvascular endothelial cell”), further emphasizing the potential loss of these cell types in the single-cell data. By contrast, we uncovered a variety of both immune and non-immune cell types overrepresented in bulk RNA-seq `ScPcP000017` (osteosarcoma; Figure S8B), which may reflect inherent challenges in dissociating bone tissue [62]. These results show that, while bulk and single-cell/nuclei expression is indeed highly correlated, cell type differences may still be present between modalities, potentially driven by cell-type-specific loss in single-cell experiments.

## Methods

### Data generation and processing

Raw data and metadata were generated and compiled by each lab and institution contributing to the Portal. Single-cell or single-nuclei libraries were generated using one of the commercially available kits from 10x Genomics. For bulk RNA-seq, RNA was collected and sequenced using either paired-end or single-end sequencing. For spatial transcriptomics, cDNA libraries were generated using the Visium kit from 10x Genomics. All libraries were processed using our open-source pipeline, `scpca-nf`, to produce summarized gene expression data. A detailed summary with the total number of samples and libraries collected for each sequencing method broken down by project is available in Table S1.

### Metadata

Submitters were required to submit the age, sex, organism, diagnosis, subdiagnosis (if applicable), disease timing (e.g., initial diagnosis) and tissue of origin for each sample. The submitted metadata was standardized across projects, including converting all ages to years, removing abbreviations used in diagnosis, subdiagnosis, or tissue of origin, and using standard values across projects as much as possible for diagnosis, subdiagnosis, disease timing, and tissue of origin. For example, all samples obtained at diagnosis were assigned the value `Initial diagnosis` for disease timing.

In an effort to ensure sample metadata for ScPCA are compatible with CZI’s CELLxGENE, ontology term identifiers were assigned to metadata categories for each sample following the guidelines present in the CELLxGENE schema [63,64], as shown in Table 1.

**Table 1:** Assignment of metadata fields to ontology terms.

| Metadata field | Ontology term description |
| --- | --- |
| Age | Ontology term obtained from HsapDv [15]. For ages 0-11 months, the HsapDv for age in months was used. For ages 12 months and greater, the HsapDv for age in years was used. |
| Sex | Ontology term obtained from PATO, either male (PATO:0000384), female (PATO:0000383), or unknown [16,17]. |
| Organism | NCBI taxonomy term for organism. All current samples available on the Portal are from Homo sapiens or NCBITaxon:9606 [18,19]. |
| Diagnosis | The most appropriate MONDO term based on the provided diagnosis [20,21]. An exact match was identified for most samples, but in a handful of cases, the most closely related term was used. |
| Tissue of origin | The most appropriate UBERON term based on the provided tissue of origin [22,23,24]. An exact match was identified for most samples, but in a handful of cases, the most closely related term was used. |
| Ethnicity (if applicable) | If the submitter provided ethnicity, the associated Hancestro term [25,26]. If ethnicity is unavailable, unknown is used. |

### Processing single-cell and single-nuclei RNA-seq data with alevin-fry

To quantify RNA-seq gene expression for each cell or nucleus in a library, `scpca-nf` uses `salmon alevin` [65] and `alevin-fry` [14] to generate a gene-by-cell counts matrix. Prior to mapping, we generated an index using transcripts from both spliced cDNA and unspliced cDNA sequences, denoted as the `splici` index [14]. The index was generated from the human genome, GRCh38, Ensembl version 104. `salmon alevin` was run using selective alignment to the `splici` index with the `--rad` option to generate a reduced alignment data (RAD) file required for input to `alevin-fry`.

The RAD file was used as input to the recommended `alevin-fry` workflow, with the following customizations. At the `generate-permit-list` step, we used the `--unfiltered-pl` option to provide a list of expected barcodes specific to the 10x kit used to generate each library. The `quant` step was run using the `cr-like-em` resolution strategy for feature quantification and UMI de-duplication.

### Post alevin-fry processing of single-cell and single-nuclei RNA-seq data

The output from running `alevin-fry` includes a gene-by-cell counts matrix, with reads from both spliced and unspliced reads for all potential cell barcodes. The gene-by-cell counts matrix is read into R to create a `SinglecellExperiment` using `fishpond::load_fry()`. The resulting `SinglecellExperiment` contains a `counts` assay with a gene-by-cell counts matrix where all spliced and unspliced reads for a given gene are totaled together. We also include a `spliced` assay that contains a gene-by-cell counts matrix with only spliced reads. These matrices include all potential cells, including empty droplets, and are provided for all Portal downloads in the unfiltered objects saved as `.rds` files with the `_unfiltered.rds` suffix.

Each droplet was tested for deviation from the ambient RNA profile using `DropletUtils::emptyDropscellRanger()` [38,66] and those with an FDR ≤ 0.01 were retained as likely cells. If a library did not have a sufficient number of droplets and `DropletUtils::emptyDropscellRanger()` failed, cells with fewer than 100 UMIs were removed. Gene expression data for any cells that remain after filtering are provided in the filtered objects saved as `.rds` files with the `_filtered.rds` suffix.

In addition to removing empty droplets, `scpca-nf` also removes cells that are likely to be compromised by damage or low-quality sequencing. `miQc` was used to calculate the posterior probability that each cell is compromised [39]. Any cells with a probability of being compromised greater than 0.75 and fewer than 200 genes detected were removed before further processing. The gene expression counts from the remaining cells were log-normalized using the deconvolution method from Lun, Bach, and Marioni [40]. `scran::modelGeneVar()` was used to model gene variance from the log-normalized counts and `scran::getTopHVGs()` was used to select the top 2000 high-variance genes. These were used as input to calculate the top 50 principal components using `scater::runPcA()`. Finally, UMAP embeddings were calculated from the principal components with `scater::runUMAP()`. The raw and log-normalized counts, list of 2000 high-variance genes, principal components, and UMAP embeddings are all stored in the processed objects saved as `.rds` files with the `_processed.rds` suffix.

### Quantifying gene expression for libraries with CITE-seq or cell hashing

All libraries with antibody-derived tags (ADTs) or hashtag oligonucleotides (HTOs) were mapped to a reference index using `salmon alevin` and quantified using `alevin-fry`. The reference indices were constructed using the `salmon index` command with the `--feature` option. References were custom-built for each ScPCA project and constructed using the submitter-provided list of ADTs or HTOs and their barcode sequences.

The ADT-by-cell or HTO-by-cell counts matrix produced by `alevin-fry` were read into R as a `SinglecellExperiment` object and saved as an alternative experiment (`altExp`) in the same `SinglecellExperiment` object with the unfiltered gene expression counts data. The `altExp` within the unfiltered object contains all identified ADTs or HTOs and all barcodes identified in the RNA-seq gene expression data. Any barcodes that only appeared in either ADT or HTO data were discarded, and cell barcodes that were only found in the gene expression data (i.e., did not appear in the ADT or HTO data) were assigned zero counts for all ADTs and HTOs. Any cells removed after filtering empty droplets were also removed from the ADT and HTO counts matrices and before creating the filtered `SinglecellExperiment` object.

### Processing ADT expression data from CITE-seq

The ADT count matrix stored in the unfiltered object was used to calculate an ambient profile with `DropletUtils::ambientProfileEmpty()`. The ambient profile was used to calculate quality-control statistics with `DropletUtils::cleanTagcounts()` for all cells remaining after removing empty droplets. Any negative or isotype controls were taken into account when calculating QC statistics. Cells with a high level of ambient contamination or negative/isotype controls were flagged as having low-quality ADT expression, but we did not remove any cells based on ADT quality from the object. The filtered and processed objects contain the results from running `DropletUtils::cleanTagcounts()`.

ADT count data were then normalized by calculating median size factors using the ambient profile with `scuttle::computeMedianFactors()`. If median-based normalization failed for any reason, ADT counts were log-transformed after adding a pseudocount of 1. Normalized counts are only available for any cells that would be retained after ADT filtering, and any cells that would be filtered out based on `DropletUtils::cleanTagcounts()` are assigned `NA`. The normalized ADT data are available in the `altExp` of the processed object. Although `scpca-nf` normalizes ADT counts, the workflow does not perform any dimensionality reduction of ADT data; only the RNA counts data are used as input for dimensionality reduction.

During conversion to AnnData objects, the modalities are exported as separate RNA (_rna.h5ad) and ADT (_adt.h5ad) objects.

### Processing HTO data from multiplexed libraries

As with ADT data, `scpca-nf` does not filter any cells based on HTO expression, and any cells removed after filtering empty droplets based on the unfiltered RNA counts matrix are also removed from the HTO counts matrix in the filtered object. `scpca-nf` does not perform any additional filtering or processing of the HTO-by-cell counts matrix, so the same filtered matrix is included in the processed object.

To identify which cells come from which sample in a multiplexed library, we applied three different demultiplexing methods: genetic demultiplexing, HTO demultiplexing using `DropletUtils::hashedDrops()`, and HTO demultiplexing using `Seurat::HTODemux()`. We do not provide separate `SinglecellExperiment` objects for each sample in a library. Each multiplexed library object contains the counts data from all samples and the results from all three demultiplexing methods to allow users to select which method(s) to use.

### Genetic demultiplexing

If all samples in a multiplexed library were also sequenced using bulk RNA-seq, we performed genetic demultiplexing using genotype data from both bulk RNA-seq and single-cell or single-nuclei RNA-seq [46]. If bulk RNA-seq was not available, no genetic demultiplexing was performed.

Bulk RNA-seq reads for each sample were mapped to a reference genome using `STAR` [67], and multiplexed single-cell or single-nuclei RNA-seq reads were mapped to the same reference genome using `STARsolo` [66]. The mapped bulk reads were used to call variants and assign genotypes with `bcftools mpileup` [68]. `cellsnp-lite` was then used to genotype single-cell data at the identified sites found in the bulk RNA-seq data [69]. Finally, `vireo` was used to identify the sample of origin [69].

### HTO demultiplexing

For all multiplexed libraries, we performed demultiplexing using `DropletUtils::hashedDrops()` and `Seurat::HTODemux()`. For both methods, we used the default parameters and only performed demultiplexing on the filtered cells present in the filtered object. The results from both these methods are available in the filtered and processed objects.

### Quantification of spatial transcriptomics data

10x Genomics’ Space Ranger [49] was used to quantify gene expression data from spatial transcriptomics libraries. `cellranger mkref` was used to create a reference index from the human genome, GRCh38, Ensembl version 104. The FASTQ files, microscopic slide image, and slide serial number were provided as input to `spaceranger count`. The raw and filtered counts matrix, slide images, and the summary report output by `spaceranger count` are included in the output from `scpca-nf`.

### Quantification of bulk RNA-seq data

`fastp` was used to trim adapters and perform quality and length filtering on all FASTQ files from bulk RNA-seq. We used a decoy-aware reference created from spliced cDNA sequences with the entire human genome sequence (GRCh38, Ensembl version 104) as the decoy [48]. The trimmed reads were then provided as input to `salmon quant` for selective alignment. In addition to using the default parameters for `salmon quant`, we applied the `--seqBias` and `--gcBias` flags to correct for sequence-specific biases due to random hexamer priming and fragment-level GC biases, respectively.

### Cell type annotation

Cell type labels determined by both `SingleR` [42] and `cellAssign` [43] were added to processed `SinglecellExperiment` objects. If cell types were obtained from the submitter of the dataset, the submitter-provided annotations were incorporated into all `SinglecellExperiment` objects (unfiltered, filtered, and processed).

To prepare the references used for assigning cell types, we developed a separate workflow, `build-celltype-index.nf`, within `scpca-nf`. For `SingleR`, we used the `BlueprintEncodeData` from the `celldex` package [51,52] to train the `SingleR` classification model with `SingleR::trainSingleR()`. In the main `scpca-nf` workflow, this model and the processed `SinglecellExperiment` object were input to `SingleR::classifySingleR()`. The `SingleR` output of cell type annotations and a score matrix for each cell and all possible cell types were added to the processed `SinglecellExperiment` object. To evaluate confidence in `SingleR` cell type assignments, we also calculated a delta median statistic for each cell by subtracting the median cell type score from the score associated with the assigned cell type [50].

For `cellAssign`, marker gene references were created using the marker gene lists available on `PanglaoDB` [53]. Organ-specific references were built using all cell types in a specified organ listed in `PanglaoDB` to accommodate all ScPCA projects encompassing a variety of disease and tissue types. If a set of disease types in a given project encompassed cells that may be present in multiple organ groups, multiple organs were combined. For example, we created a reference containing bone, connective tissue, smooth muscle, and immune cells for sarcomas that appear in bone or soft tissue.

Given the processed `SinglecellExperiment` object and organ-specific reference, `scvi.external.cellAssign()` was used in the main `scpca-nf` workflow to train the model and predict the assigned cell type. For each cell, `cellAssign` calculates a probability of assignment to each cell type in the reference. The probability matrix and a prediction based on the most probable cell type were added as cell type annotations to the processed `SinglecellExperiment` object.

### Assigning consensus cell types

Cell type labels obtained from `SingleR` and `cellAssign` were then used to assign an ontology-aware consensus cell type label. We first assigned each of the cell types present in the `PanglaoDB` [53] reference used with `cellAssign` to an appropriate Cell Ontology term [70]. For cell types available in the `BlueprintEncodeData` reference used with `SingleR`, we used the provided Cell Ontology terms.

We then created a reference table containing all possible combinations of cell types assigned using `SingleR` and `cellAssign` and identified the latest common ancestor (LCA) using `ontoProc::findcommonAncestors()` [71] between the two cell type terms. The LCA was then used as the consensus cell type label if the following criteria were met, otherwise no consensus cell type was assigned:

1. The terms shared only one distinct LCA. There was one exception to this rule: If the terms shared two LCAs and one was `hematopoietic precursor cell`, then `hematopoietic precursor cell` was used as the consensus label.
2. The LCA had fewer than 170 descendants, or was either `neuron` or `epithelial cell`.

If the LCA was one of the following non-specific LCA terms, no consensus cell type was assigned: bone cell, lining cell, blood cell, progenitor cell, and supporting cell.

The consensus cell type assignments, including both the Cell Ontology term and the associated human-readable name, are available in processed object files on the Portal.

Consensus cell type assignments were evaluated by looking at marker gene expression in a set of cell-type specific marker genes. Marker genes were obtained from the list of Human cell markers on `cellMarker2.0` [72]. We considered only those that are specific to a single cell type, with the exception of hematopoietic precursor cells, which express genes found in other, more differentiated immune cells.

### Generating merged data

Merged objects are created with the `merge.nf` workflow within `scpca-nf`. This workflow takes as input the processed `SinglecellExperiment` objects in a given ScPCA project output by `scpca-nf` and creates a single merged `SinglecellExperiment` object containing gene expression data and metadata from all libraries in that project. The merged object includes both raw and normalized counts for all cells from all libraries. Because the same reference index was used to quantify all single-cell and single-nuclei RNA-seq data, the set of genes is the same in the merged object and the individual objects. Library-, cell- and gene-specific metadata from each of the processed `SinglecellExperiment` objects are also combined and stored in the merged object. The `merge.nf` workflow does not perform batch correction or integration, so the counts in the merged object are not batch-corrected.

The top 2000 shared high-variance genes are identified from the merged counts matrix by modeling variance using `scran::modelGeneVar()` and specifying library IDs for the `block` argument. These genes are used to calculate library-aware principal components with `batchelor::multiBatchPcA()` [73]. The top 50 principal components were selected and used to calculate UMAP embeddings for the merged object.

If any libraries included in the ScPCA project contain additional ADT data, the raw and normalized ADT data are also merged and stored in the `altExp` slot of the merged `SinglecellExperiment` object. If the merged object contains an `altExp` with merged ADT data, two `AnnData` objects are exported to create separate RNA (`_rna.h5ad`) and ADT (`_adt.h5ad`) objects.

If any libraries in the ScPCA project are multiplexed and contain HTO data, no merged object is created due to potential ambiguity in identifying samples across multiplexed libraries. Merged objects were not created for projects with more than 100 samples because of the computational resources required to work with them.

### Converting SingleCellExperiment objects to AnnData objects

`zellkonverter::writeH5AD()` [74] was used to convert `SinglecellExperiment` objects to `AnnData` format and export the objects as `.h5ad` files. For any `SinglecellExperiment` objects containing an `altExp` (e.g., ADT data), the RNA and ADT data were exported and saved separately as RNA (`_rna.h5ad`) and ADT (`_adt.h5ad`) files. Multiplexed libraries were not converted to `AnnData` objects, due to the potential for ambiguity in sample origin assignments.

All merged `SinglecellExperiment` objects were converted to `AnnData` objects and saved as `.h5ad` files. If a merged `SinglecellExperiment` object contained any ADT data, the RNA and ADT data were exported and saved separately as RNA (`_rna.h5ad`) and ADT (`_adt.h5ad`) objects. In contrast, if a merged `SinglecellExperiment` object contained HTO data due to the presence of any multiplexed libraries in the merged object, the HTO data was removed from the `SinglecellExperiment` object and not included in the exported `AnnData` object.

### Analysis of bulk RNA-seq data

#### Data preparation

We identified solid tumor samples with both bulk and single-cell (or single-nuclei) RNA-seq data in the ScPCA Portal for analysis, with multiplexed samples excluded (N=105). We removed low-quality samples based on visual inspection of quality control reports (N=8), leaving a total of 97 samples across five ScPCA projects for analysis.

For each project, we transformed and normalized bulk counts matrices for all samples using `DESeq2::rlog()` [75]. We obtained pseudobulk counts by summing raw single-cell counts for each sample, and similarly transformed each project’s resulting counts matrix with `DESeq2::rlog()`. We filtered out genes which were not observed in either the bulk or pseudobulk raw counts matrices before subsequent analysis. For each project, we then used the `lme4` [76] R package to construct a linear model predicting bulk from pseudobulk counts considering a random effect for sample id: `bulk ∼ pseudobulk + (1|sample_id)`.

#### Overrepresentation analysis

We next asked whether certain cell types might be overrepresented in one modality compared to the other. For this, we first identified cell types of interest as the set of all possible consensus cell types for each project. We then created a gene set for each consensus cell type using the project’s `cellAssign` marker gene reference. Because a consensus cell type can encompass multiple cell types in the marker gene reference, we defined each consensus cell type’s gene set as the union of all marker genes for each of its constituent reference cell types.

For input to the overrepresentation analysis, we summarized model residuals within each project by taking the median residual for each gene across samples and then transformed these summarized residuals into Z-scores. We identified outlier genes as those with Z-scores greater than 2.5 (positive outliers) or less than −2.5 (negative outliers). In this case, positive outliers represent genes with comparatively higher expression in the bulk modality, and negative outliers represent genes with comparatively higher expression in the single-cell modality.

For each consensus cell type gene set, we calculated two odds ratios representing whether genes were overrepresented in the positive outliers (enriched in bulk) or negative outliers (enriched in pseudobulk). We calculated P-values for both the bulk and pseudobulk enrichment directions via permutation testing with 10,000 replicates. We defined gene sets with significant overrepresentation as those with a false-discovery-rate-corrected P-value ≤ 0.05 [77].

## Code and data availability

All summarized gene expression data and de-identified metadata are available for download on the ScPCA Portal, https://scpca.alexslemonade.org/.

Documentation for the Portal can be found at https://scpca.readthedocs.io.

All original code was developed within the following repositories and is publicly available as follows:

- The scpca-nf workflow used to process all samples available on the Portal can be found at https://github.com/AlexsLemonade/scpca-nf.
- The Single-cell Pediatric Cancer Atlas Portal code can be found at https://github.com/AlexsLemonade/scpca-portal
- Benchmarking of tools used to build scpca-nf can be found at https://github.com/AlexsLemonade/alsf-scpca/tree/main/analysis and https://github.com/AlexsLemonade/sc-data-integration/tree/main/celltype_annotation.
- All code for creating the reference files used for consensus cell type assignment can be found at https://github.com/AlexsLemonade/OpenScPCA-analysis/tree/main/analyses/cell-type-consensus.
- All code for the underlying figures and analyses can be found at https://github.com/AlexsLemonade/scpca-paper-figures.
- The manuscript can be found at https://github.com/AlexsLemonade/ScPCA-manuscript.

## Discussion

The ScPCA Portal is a downloadable collection of uniformly processed, summarized single-cell and single-nuclei RNA-seq data and de-identified metadata from pediatric tumor samples. The Portal includes over 700 samples from 55 tumor types, making this the most comprehensive collection of publicly available single-cell RNA-seq datasets from pediatric tumor samples to our knowledge.

Summarized data are available at three different processing stages: unfiltered, filtered, or processed objects. Users can choose to start from a processed object or perform their own processing, such as filtering and normalization. Processed objects containing normalized gene expression data, reduced dimensionality results from PCA and UMAP, and cell type annotations. Standardized metadata, containing human-readable values for all fields and ontology term identifiers for a subset of metadata fields, is included in a separate metadata file and the data objects for all samples. Every library includes a quality control report, which lets users assess data quality and identify low-quality libraries that they may wish to exclude from further downstream analyses. The availability of processed results and metadata saves time and effort for researchers, allowing them to move directly to downstream analyses, such as identifying marker genes or exploring genes of interest.

Data on the Portal is available as either `SinglecellExperiment` or `AnnData` objects so that users can work in R or Python with the downloaded data using common analysis systems such as `Bioconductor` or `Scanpy`, depending on their preference. Providing data as `AnnData` objects also means users can easily integrate ScPCA data with data and tools available on other platforms. In particular, the format of the provided `AnnData` objects was designed to be mostly compliant with the requirements of CZI CELLxGENE [5,6,78], but these objects can also be used with UCSC Cell Browser [79,80] or Kana [81,82]. Additionally, users can download a merged `SinglecellExperiment` or `AnnData` object containing all gene expression data and metadata from all samples in a project, which supports multi-sample analyses such as differential gene expression or gene set enrichment.

To provide users with cell type annotations, we used two automated methods, `SingleR` and `cellAssign`, which use publicly available references. We then used the correspondence between methods to derive ontology-aware consensus cell type labels. A limitation of our annotation approach is that the references we used do not contain tumor cells; therefore, tumor cells are likely poorly assigned. However, the consensus cell type labels provide a consistent labeling scheme across samples and may be beneficial for annotating populations of normal cells that may be present in tumor samples.

Many samples on the Portal have additional sequencing data, including corresponding ADT data from CITE-seq, cell hashing data, bulk RNA-seq, or spatial transcriptomics. This enables users to gather more information about a single sample than they could from single-cell/nuclei RNA-seq alone.

Samples with CITE-seq have additional information about cell-surface protein expression in individual cells, which can help determine cell types and correlate RNA to protein expression [34]. Spatial transcriptomics data on the Portal are not single-cell resolution, making it hard to identify cell types and spatial patterns from the spatial data alone. By providing matching single-cell RNA-seq, users can implement analysis tools, like those that use single-cell RNA-seq to deconvolute spatial data, to gain more insights about the spatial data [83].

Similarly, users can gain more insight from bulk RNA-seq data available on the Portal by integrating with single-cell RNA-seq data from the same sample [84,85]. The single-cell RNA-seq data available on the Portal can also be used to deconvolute existing bulk RNA-seq datasets, allowing researchers to infer the abundance of different cell types or cell states in bulk RNA-seq data. Our analysis of this data showed that while expression is generally consistent between matched bulk RNA-seq and single-cell/nuclei libraries in the Portal (Figure 6A, Figure S8A), there are potential differences in cell type composition between modalities that may reflect technological differences in sample and library preparation. The ScPCA Portal enables multimodal comparisons that reveal biological and/or technical signals that would otherwise not be apparent from one sequencing modality alone.

We also introduced our open-source and efficient workflow for uniformly processing datasets available on the Portal, `scpca-nf`, which is available to the entire research community. In one command, `scpca-nf` can process raw data from various sequencing types, turning FASTQ files into processed `SinglecellExperiment` or `AnnData` objects ready for downstream analyses. Using Nextflow as the framework for `scpca-nf` with Docker images for each process makes the workflow modular and portable. This makes it easy to add support for more modalities in the future, such as single-cell ATAC-seq, and allows others to run the workflow on their samples in their computing environment, maintaining the security of protected raw data. Processed output from running `scpca-nf` on samples from pediatric tumors, cell lines, or other model organisms is eligible for submission to the ScPCA Portal, enabling us to continue to grow the Portal.

Data available on the ScPCA Portal can further be used for external data analyses, for example, to support re-analyzing any existing pediatric cancer datasets with bulk RNA-seq, such as the Pediatric Brain Tumor Atlas [86,87]. This allows researchers to glean more insight from previously published data without obtaining additional samples, saving time and money and advancing biomedical research.

## Acknowledgments

We thank the data generators and submitters of the Single-cell Pediatric Cancer Atlas. We also thank Anna Greene for her role in constructing the Single-cell Pediatric Cancer Atlas funding opportunity.

This work was funded through the Alex’s Lemonade Stand Foundation Childhood Cancer Data Lab and Childhood Cancer Data Lab Postdoctoral Fellowship (SMF).

## Author Contributions

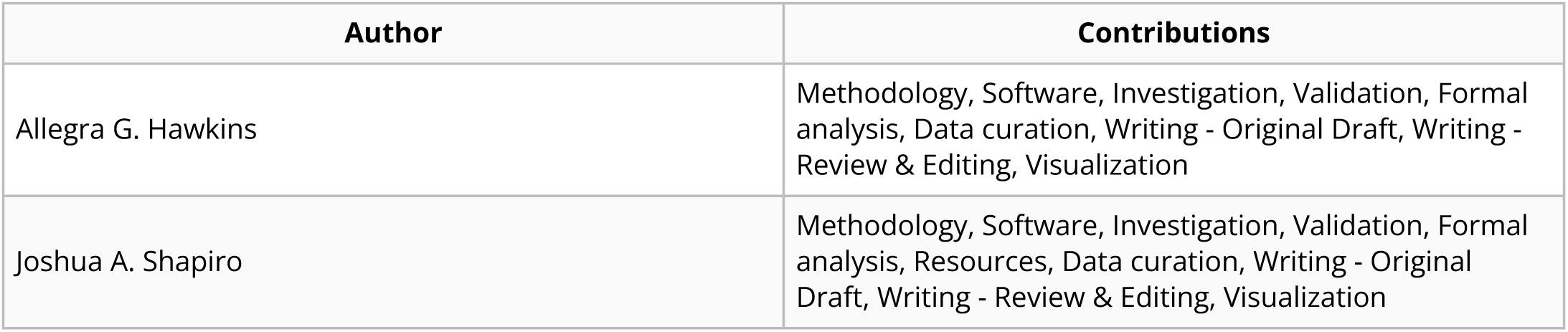
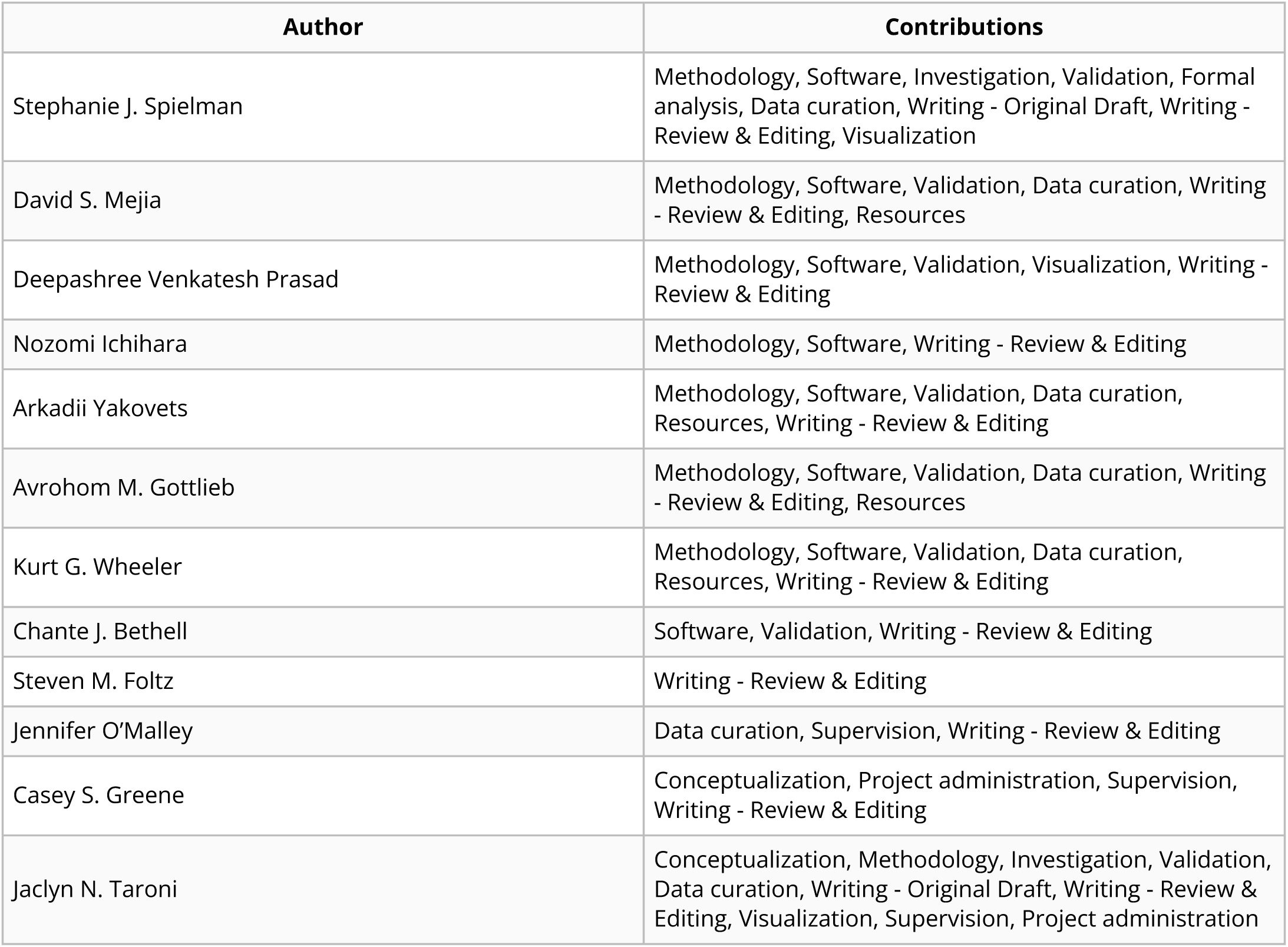

## Declarations of Interest

AGH, JAS, SJS, DSM, DVP, NI, AY, AMG, KGW, CJB, JO, and JNT are or were employees of Alex’s Lemonade Stand Foundation, a sponsor of this research.

## Supplementary Figures and Tables

**Table S1. Overview of ScPCA Portal Datasets.** This table provides descriptions and sample and library counts for each project in the ScPCA Portal.

`scpca_project_id` : ScPCA project unique identifier. `Diagnosis group` : Diagnosis group as shown in Figure 1. `Diagnoses` : Full set of diagnoses for all samples associated with the project. `Total number of samples (S)` : Number of samples associated with the project. `Total number of libraries (L)` : Number of libraries associated with the project. Due to additional sequencing modalities and/or multiplexing, projects may have more libraries than samples. All remaining columns give the number of libraries (as designated with `(L)`) with the given suspension type, 10x kit version, or additional modality.

**Table S2. Summary of references used for cell type annotation with `cellAssign`.** This table provides a summary of the references used for assigning cell types for ScPCA projects using `cellAssign`. All references were built using all cell types from a specified set of organs present in `PanglaoDB`’s marker gene list.

`scpca_project_id` : ScPCA project unique identifier. `Diagnoses` : Full set of diagnoses for all samples associated with the project. `ScPcA reference name` : Name used to describe the custom reference. `PanglaoDB organs included in reference` : A list of all organs included in the reference with names of organs corresponding to organs listed in `PanglaoDB`. The reference includes marker genes for all cell types present in each organ.

**Figure S1:**
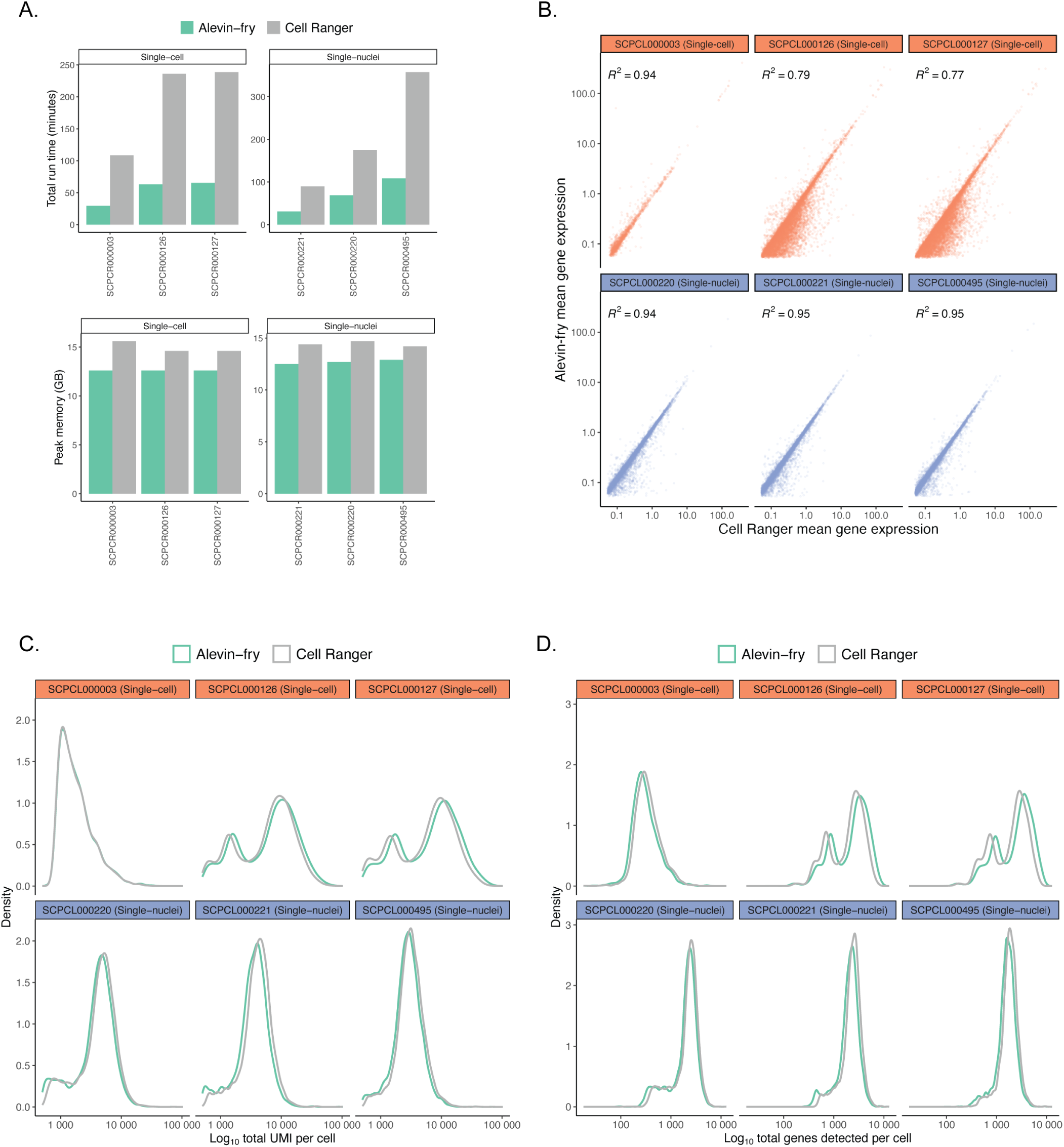
Results from benchmarking alevin-fry and cellRanger performance. Each panel compares metrics for six representative ScPCA libraries, including three single-cell and three single-nuclei suspensions, obtained from processing libraries with both `alevin-fry` and `cellRanger`. A. Runtime in minutes (top row) and peak memory in GB (bottom row) for six ScPCA libraries processed with `alevin-fry` and `cellRanger`. Processing with `alevin-fry` was consistently faster and more memory-efficient compared to processing with `cellRanger`. Panels B-D show only cells present in both the `alevin-fry` and `cellRanger` output. B. Comparison of mean gene expression values for six ScPCA libraries processed with `alevin-fry` and `cellRanger`, shown on a log-scale. Each point is a gene, and only genes detected in at least 5 cells are shown. *R*^2^ values shown in the top left corner of each panel reflect broad agreement in mean gene expression values between platforms. C. Comparison of log total UMI counts for six ScPCA libraries processed with `alevin-fry` and `cellRanger`. Distributions reflect broad agreement in the total UMI count per cell between platforms, although `alevin-fry` returned slightly higher values for certain single-cell libraries. D. Comparison of log total genes detected per cell for six ScPCA libraries processed with `alevin-fry` and `cellRanger`. Distributions reflect broad agreement between platforms in the total number of genes detected per cell between platforms, although `alevin-fry` returned slightly higher values for certain single-cell libraries.

**Figure S2:**
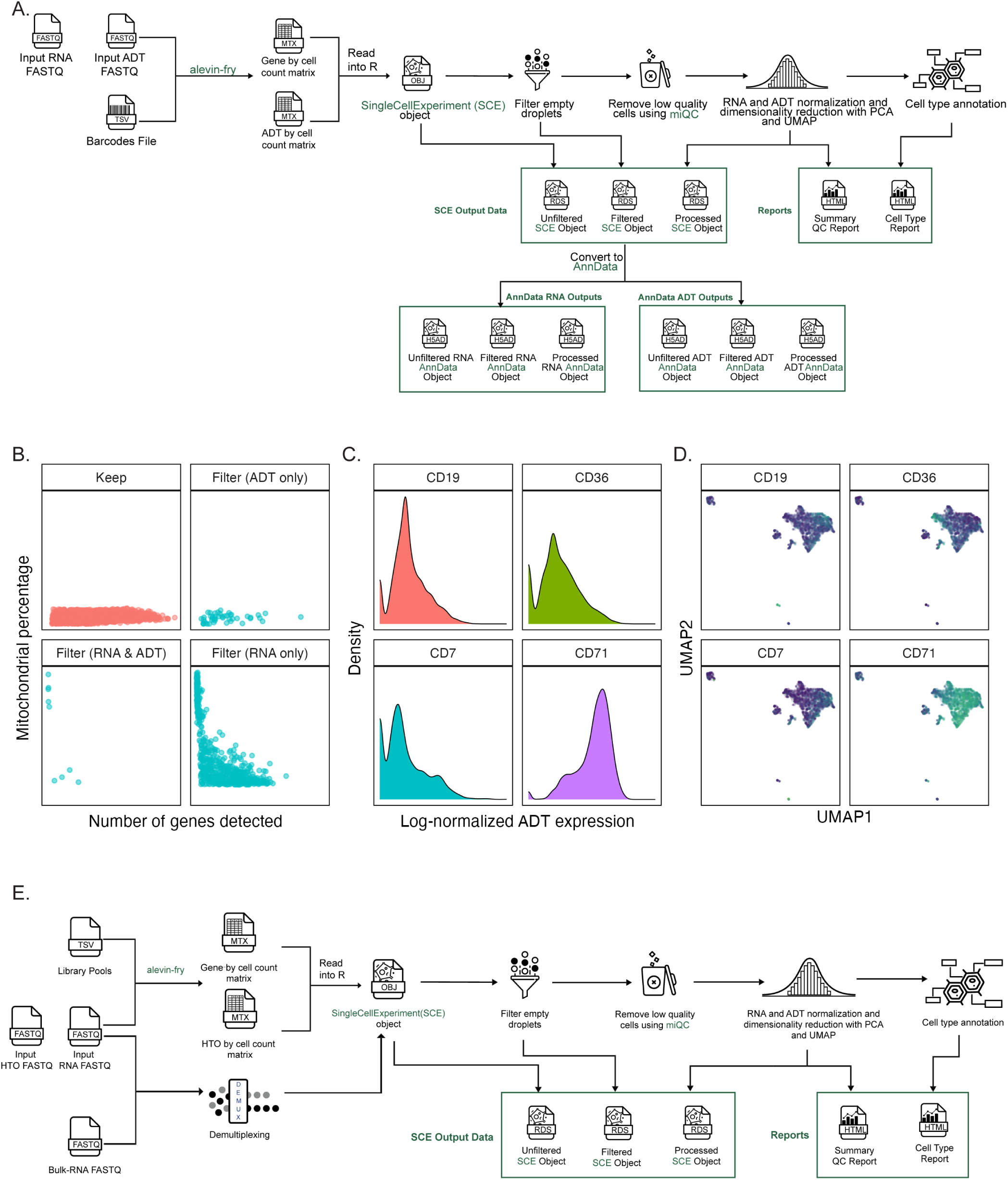
Processing additional single-cell modalities in scpca-nf. A. Overview of the `scpca-nf` workflow for processing libraries with CITE-seq or antibody-derived tag (ADT) data. The workflow mirrors that shown in Figure 2A with several differences accounting for the presence of ADT data. First, both an RNA and ADT FASTQ file are required as input to `alevin-fry`, along with a TSV file containing information about ADT barcodes. The gene-by-cell and ADT-by-cell count matrices are produced and read into `R` to create a `SinglecellExperiment` (SCE) object. Second, during post-processing, statistics are calculated to filter cells based on ADT counts, but the filter is not applied. ADT counts are also normalized and included in the `Processed ScE Object`. Third, the summary QC report will include a `cITE-seq` section with additional information about ADT-level processing. Fourth, the workflow exports `ScE` objects containing both RNA and ADT results, while separate `AnnData` objects for RNA and ADT are exported. Panels B-D show example figures that appear in the CITE-seq section of the summary QC report, shown here for `ScPcL000260`. B. The percent of mitochondrial reads in each cell against the number of genes detected in each cell. The panel labeled “Keep” displays cells that are retained based on both RNA and ADT counts. The panel labeled “Filter (ADT only)” displays cells that are filtered based on only ADT counts. The panel labeled “Filter (RNA only)” displays cells that are filtered based on only RNA counts. The panel labeled “Filter (RNA & ADT)” panel displays cells that are filtered based on both RNA and ADT counts. C. Density plots of the log-normalized ADT counts shown for the four most variable ADTs in the library. D. UMAP embeddings of log-normalized RNA expression values where each cell is colored by the expression of the given highly-variable ADT. E. Overview of the `scpca-nf` workflow for multiplexed libraries. The workflow mirrors that shown in Figure 2A with several differences accounting for the presence of multiplexed data. First, both an RNA and HTO FASTQ file are required as input to `alevin-fry`, along with a TSV file providing information about library pools. The gene-by-cell and HTO-by-cell count matrices are produced and read into `R` to create a `SinglecellExperiment` (SCE) object. Second, in parallel, the RNA FASTQ file, the HTO FASTQ file, and, if available, a corresponding Bulk RNA FASTQ file for each sample present in the multiplexed library are provided to a demultiplexing subprocess. The workflow calculates demultiplexing results based on HTO counts, as well as genetic demultiplexing results if the library has corresponding bulk RNA FASTQ files. Demultiplexing results are stored in all exported `ScE` objects (`Unfiltered`, `Filtered`, and `Processed`), but libraries themselves are not demultiplexed. Third, only `ScE` files are provided for multiplexed libraries; no corresponding `AnnData` files are provided.

**Figure S3:**
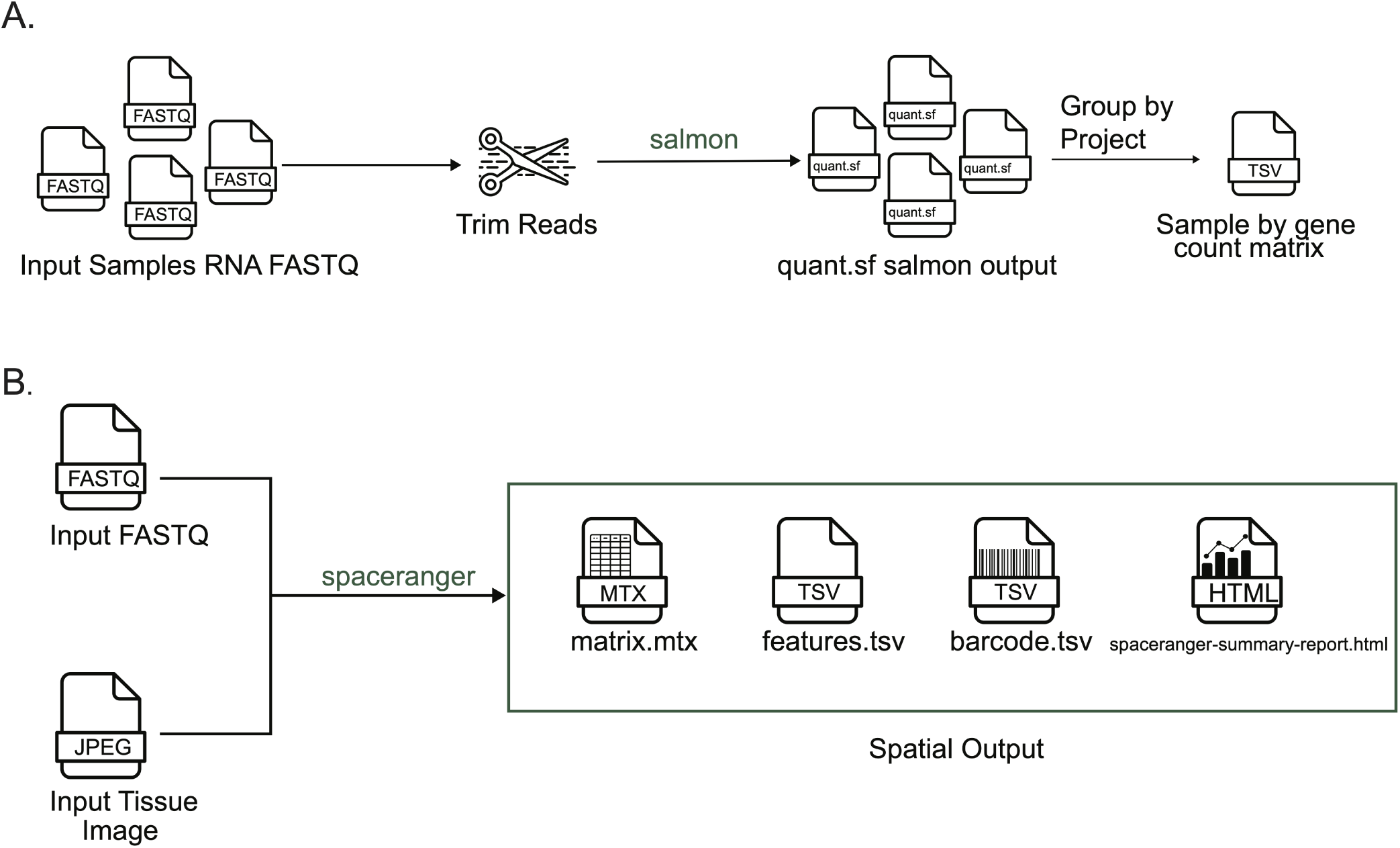
Processing other sequencing modalities with scpca-nf. A. Overview of the bulk RNA-Seq workflow. A set of FASTQ files from libraries sequenced with bulk RNA-seq are provided as input. Reads are trimmed using `fastp`, and `salmon` is used to map reads and quantify counts. The quantified gene expression files output from `salmon` are then grouped by ScPCA Project ID, and a sample-by-gene count matrix is exported for each Project in TSV format. B. Overview of the spatial transcriptomics workflow. The FASTQ file and tissue image for a given library are provided as input to `spaceranger`. The workflow directly returns the results from running `spaceranger` without any further processing.

**Figure S4:**
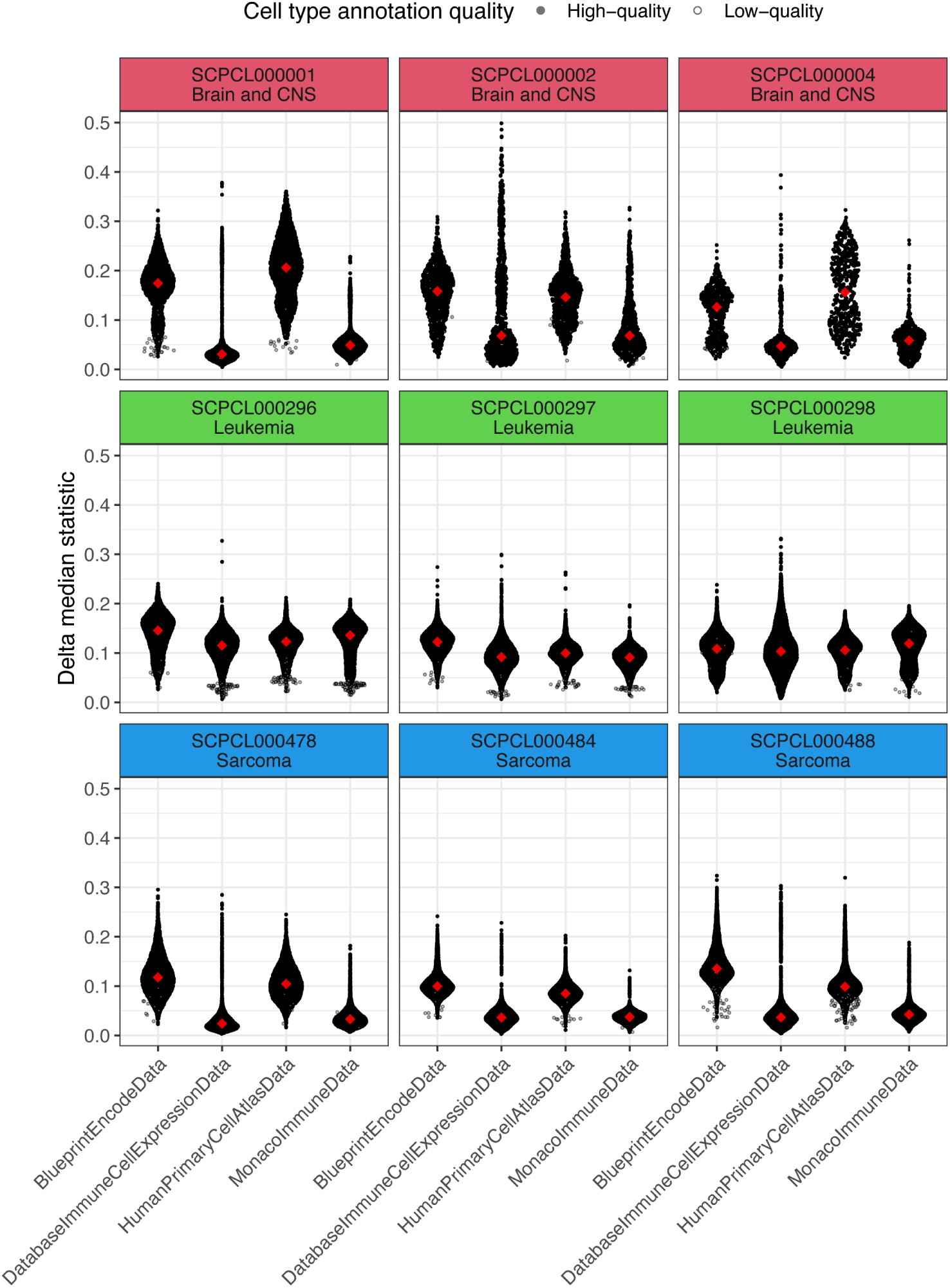
Evaluation of references available in the celldex package for use with SingleR. `SingleR` was used to annotate ScPCA libraries using four different human-specific references from the `celldex` package. Libraries represent three different diagnosis groups in the ScPCA Portal - Brain and CNS, Leukemia, and Sarcoma - as indicated in the labels for the individual panels. The distribution of the delta median statistic, calculated for each cell by subtracting the median delta score from the score of the annotated cell type label, is shown on the y-axis, while the `celldex` reference used is shown on the x-axis. Higher values indicate a higher quality cell type annotation, although there is no absolute scale for these values. Each black point represents a cell, where closed circles denote cells with high-quality annotations and open circles denote cells with low-quality annotations, as assessed by `SingleR`. Red diamonds represent the median delta median score for all cells with high-quality annotations in that library.

**Figure S5:**
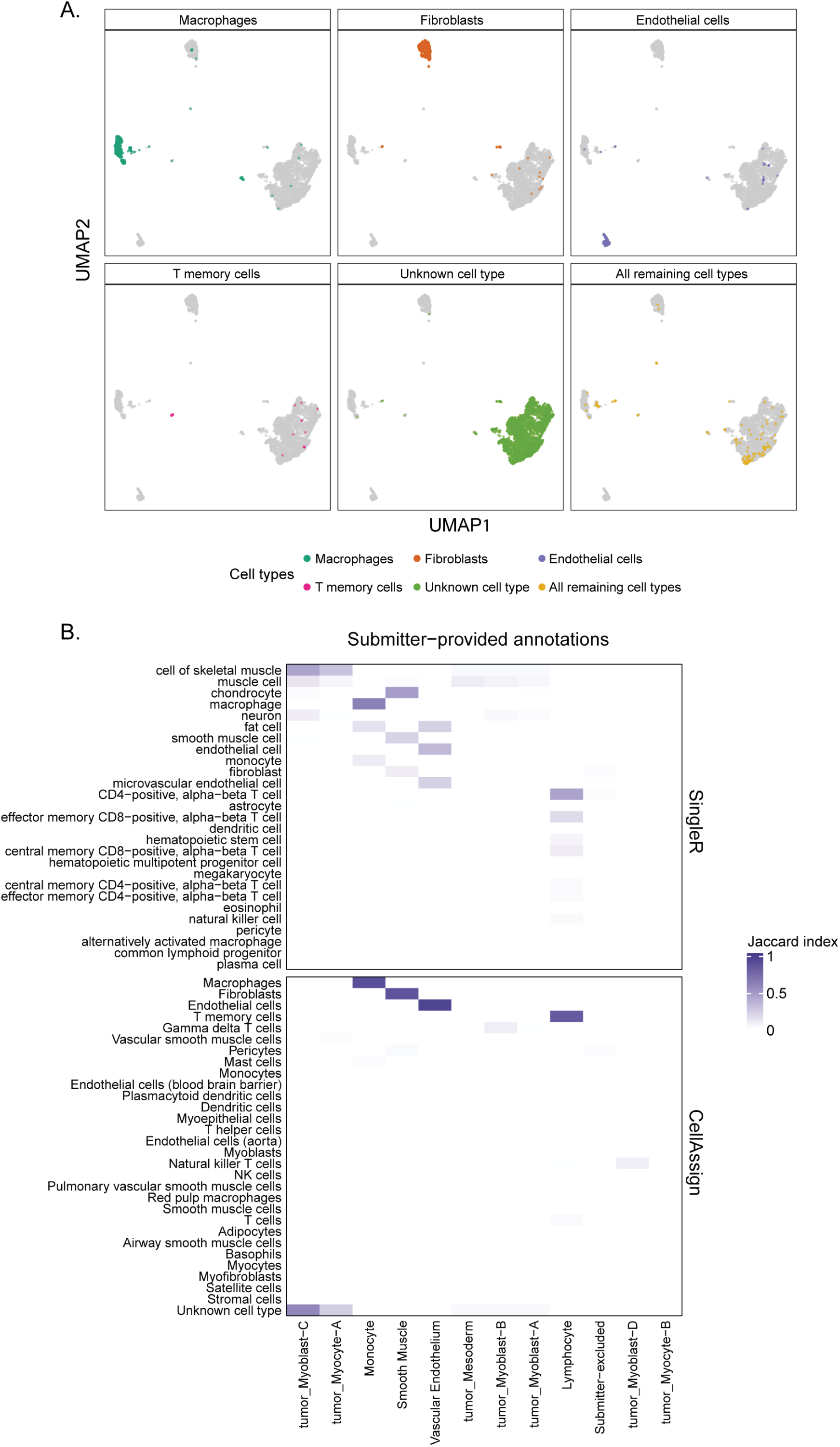
Cell type annotation with cellAssign. Both plots in this figure are examples of plots that display results from annotating cells with `cellAssign` that can be found in the cell type summary report, shown here for library `ScPcL000468` [27]. A. A grid of UMAPs is shown for each cell type annotated using `cellAssign`, with the cell type of interest shown in color and all other cells belonging to other cell types shown in gray. The top four cell types with the greatest number of assigned cells are shown, while all other cells are grouped together and labeled with `All remaining cell types`. Any cells that are unable to be assigned by `cellAssign` are labeled with `Unknown cell type`. B. This example heatmap from the cell type summary report compares submitter-provided annotations to annotations with `SingleR` and `cellAssign`. This heatmap is only shown in the cell type summary report if submitters provided cell type annotations. Heatmap cells are colored by the Jaccard similarity index. A value of 1 means that there is complete overlap between which cells are annotated with the two labels being compared, and a value of 0 means that there is no overlap between which cells are annotated with the two labels being compared.

**Figure S6:**
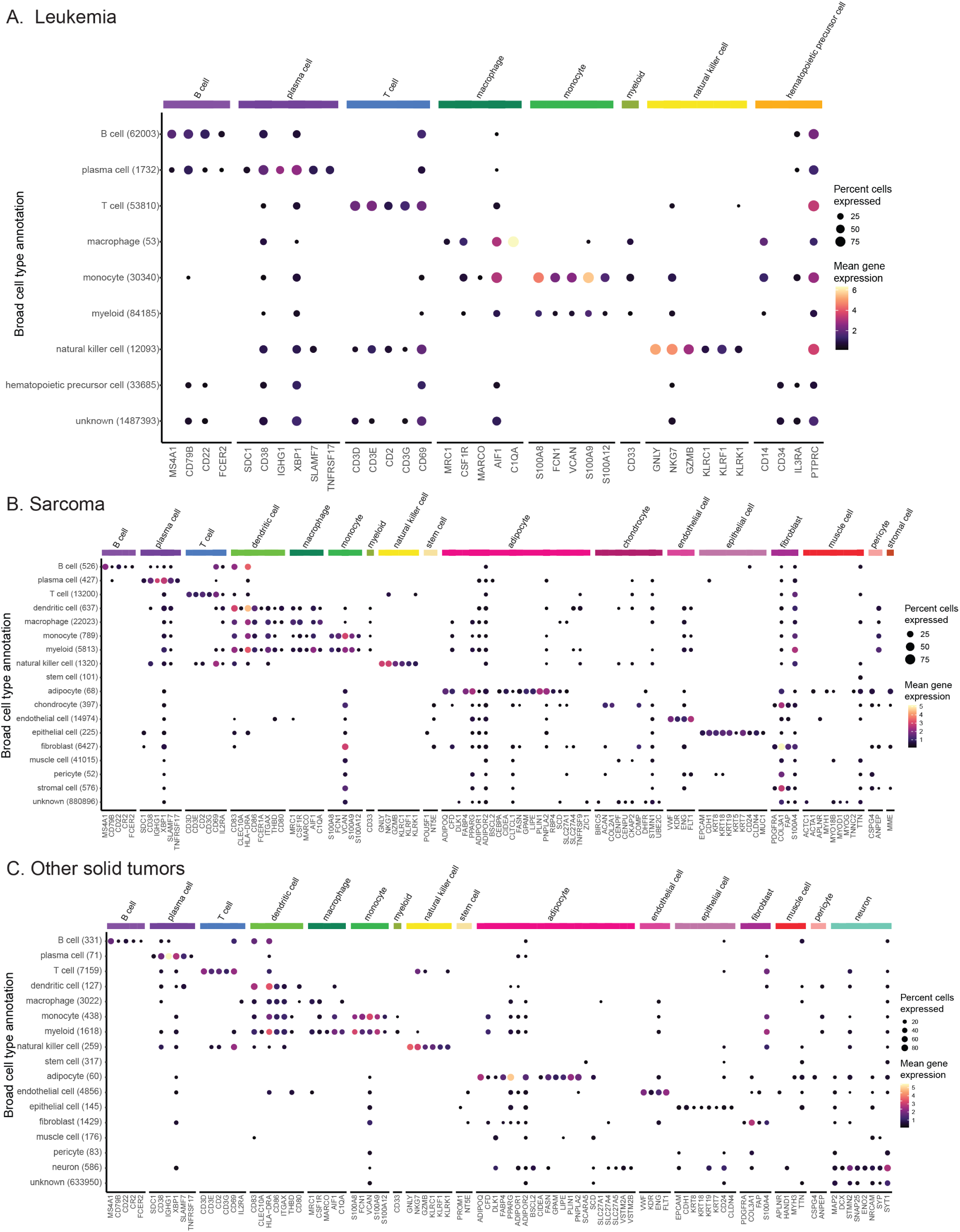
Consensus cell type annotation gene expression in other diagnosis groups. Dot plots showing expression of cell-type-specific marker genes across all libraries from Leukemia (A), Sarcoma (B), and Other solid tumors (C) diagnosis groups. Expression is shown for each broad cell type annotation, where each broad cell type annotation is a collection of similar consensus cell type annotations. The y-axis displays the broad consensus cell type observed across libraries, with the total number of cells indicated in parentheses. The x-axis displays marker genes, determined by `cellMarker2.0` [72], used for consensus cell type validation for each cell type shown along the top annotation bar. Dots are colored by mean gene expression across libraries and sized proportionally to the percent of libraries they are observed in, out of all cells with the same broad cell type annotation in the given diagnosis.

**Figure S7:**
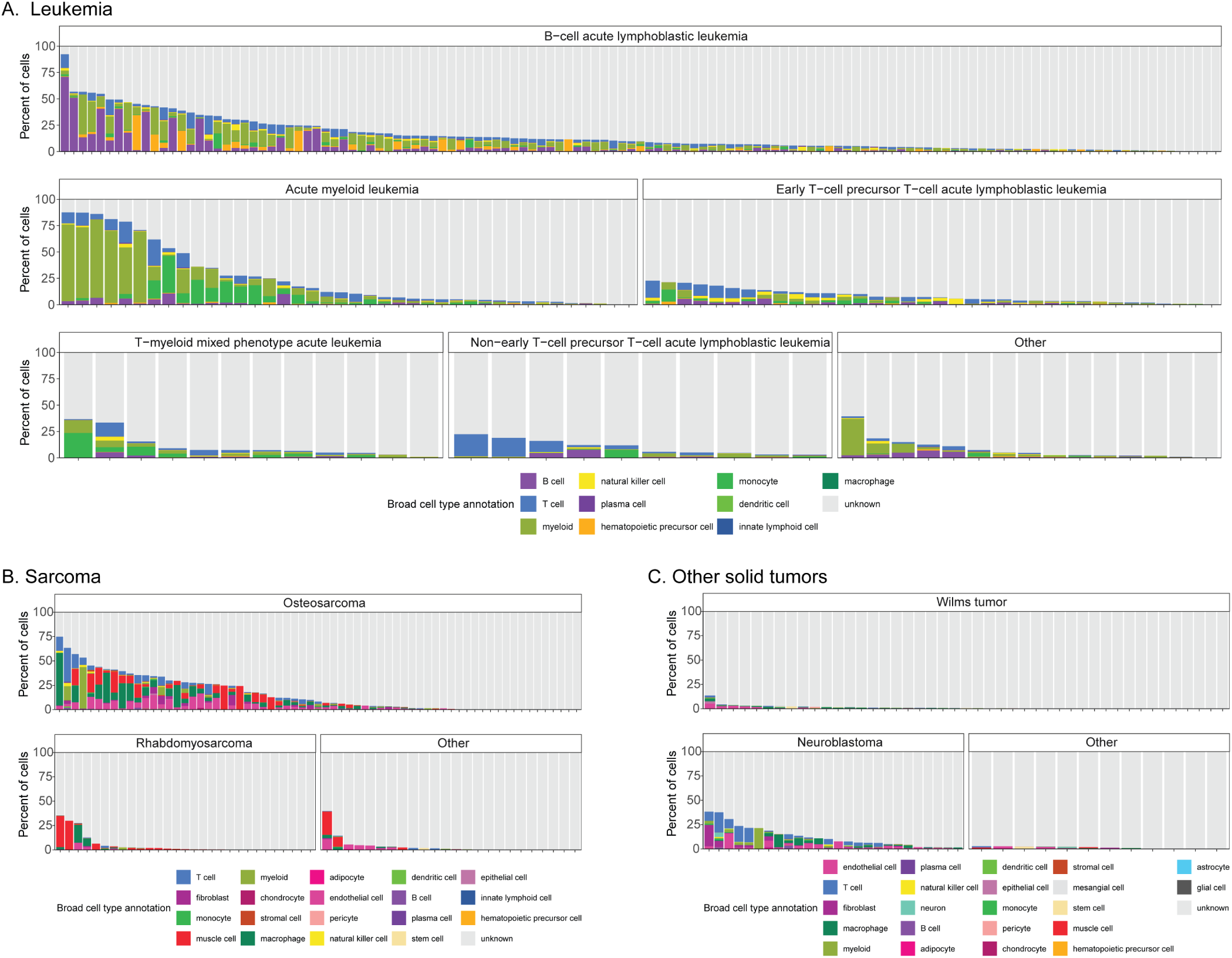
*Consensus cell type annotation distributions in other diagnosis groups. Barplots of the percentage of cells annotated as each broad consensus cell type annotation across all libraries from Leukemia (A), Sarcoma (B), and Other solid tumors (C) diagnosis groups. Within each panel, libraries are shown grouped by diagnosis. Each column represents the distribution of cell types within a single library.

**Figure S8:**
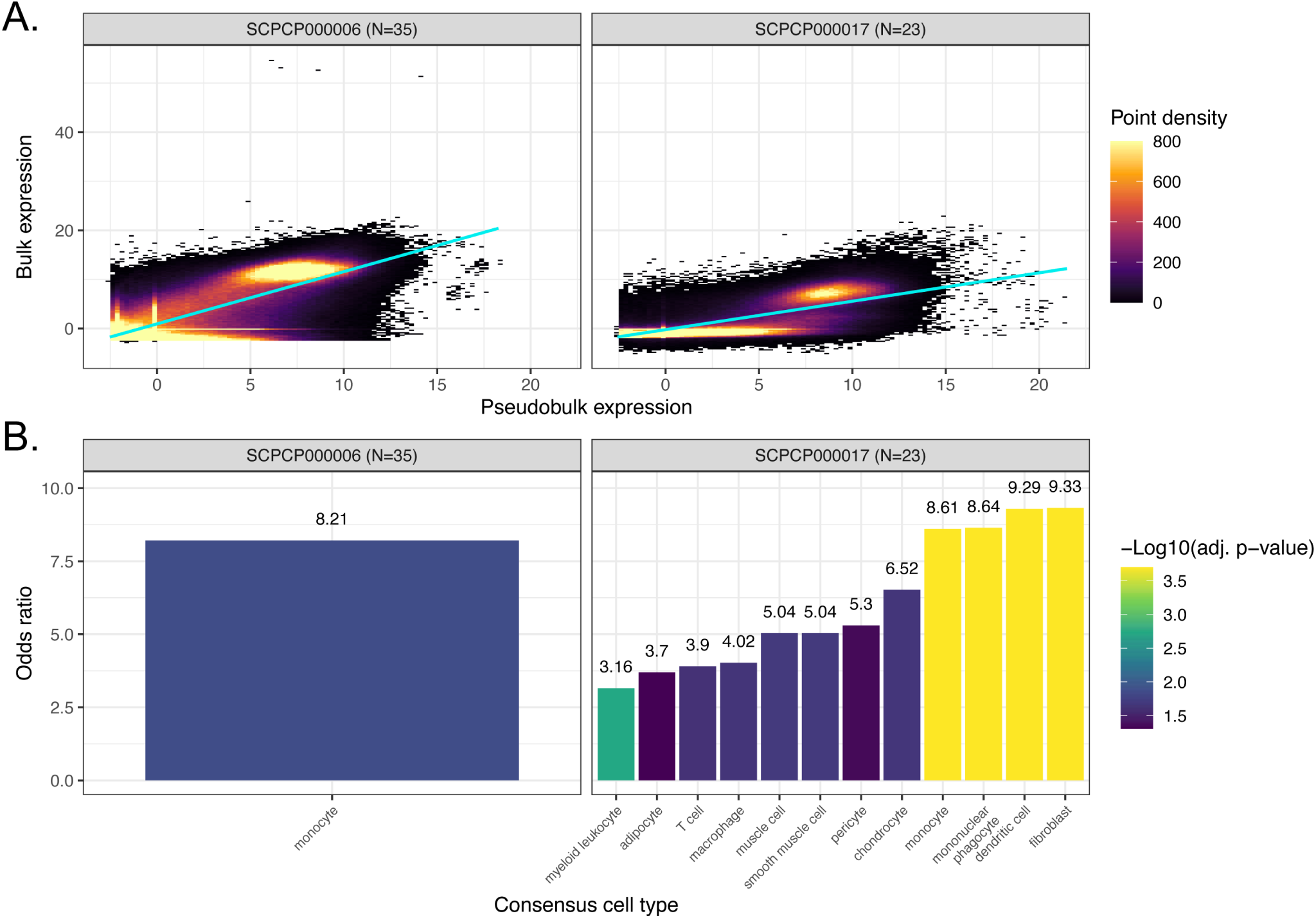
Comparison of bulk and pseudobulk modalities for additional projects. A. Scatter plots colored by point density of `DESeq2` -transformed and normalized bulk RNA-seq expression compared to pseudobulk expression from single-nuclei RNA-seq. Projects with RNA-seq for both bulk and single-cell/nuclei modalities that are not displayed in Figure 6A are shown. All samples shown here are single-nuclei, and the number of samples considered per project is shown in parentheses. The regression line is also shown for each project. B. Odds ratios from overrepresentation analysis for the same samples shown in panel A, colored by FDR-corrected significance. Each odds ratio represents the odds that marker genes for the given cell type were overrepresented in the bulk modality, relative to other genes. 31 consensus cell types were evaluated for project `ScPcP000006`, and 37 consensus cell types were evaluated for project `ScPcP000017`.

## Notes

https://scpca.alexslemonade.org/

### Summary of Updates

New Figures 5, 6, S6, S7, and S8 cover consensus cell type calls and a comparison between bulk and pseudobulk modalities; methods are updated to account for new analyses in figures as mentioned; an additional author is included.

